# The Molecular Architecture of the Nuclear Basket

**DOI:** 10.1101/2024.03.27.587068

**Authors:** Digvijay Singh, Neelesh Soni, Joshua Hutchings, Ignacia Echeverria, Farhaz Shaikh, Madeleine Duquette, Sergey Suslov, Zhixun Li, Trevor van Eeuwen, Kelly Molloy, Yi Shi, Junjie Wang, Qiang Guo, Brian T. Chait, Javier Fernandez-Martinez, Michael P. Rout, Andrej Sali, Elizabeth Villa

**Author notes:** Department of Pharmacological Sciences, Icahn School of Medicine at Mount Sinai, 1425 Madison Avenue, 16-78B, New York, NY 10029, USA. These authors contributed equally to this work (Digvijay Singh, Neelesh Soni, Joshua Hutchings). Digvijay Singh: dgvjayS. Neelesh Soni: neeleshsoni03. Joshua Hutchings: Josh-Hutchings94. Ignacia Echeverria: ig_ech. Trevor van Eeuwen: _themovingvan_. Michael P. Rout : Rout-Lab_RU. Andrej Sali : salilab_ucsf. Elizabeth Villa : TheVillaLab.

## Abstract

The nuclear pore complex (NPC) is the sole mediator of nucle-ocytoplasmic transport. Despite great advances in understanding its conserved core architecture, the peripheral regions can exhibit considerable variation within and between species. One such structure is the cage-like nuclear basket. Despite its crucial roles in mRNA surveillance and chromatin organization, an architectural understanding has remained elusive. Using in-cell cryo-electron tomography and subtomogram analysis, we explored the NPC’s structural variations and the nuclear basket across fungi (yeast; *S. cerevisiae*), mammals (mouse; *M. musculus*), and protozoa (*T. gondii*). Using integrative structural modeling, we computed a model of the basket in yeast and mammals that revealed how a hub of Nups in the nuclear ring binds to basket-forming Mlp/Tpr proteins: the coiled-coil domains of Mlp/Tpr form the struts of the basket, while their unstructured termini constitute the basket distal densities, which potentially serve as a docking site for mRNA preprocessing before nucleocytoplasmic transport

## Introduction

The nuclear pore complex (NPC) is a massive macromolecular assembly in the nuclear envelope (NE), responsible for nucleocytoplasmic transport (1–4). It comprises hundreds of proteins of more than 30 different types, known as Nucleoporins (Nups). These Nups, present in copies ranging from 8 to 48, assemble into multiple rings stacked along the nuclear envelope (1–10). These include outer rings on the nuclear and cytoplasmic sides (nuclear ring, NR; and cytoplasmic ring, CR) and an inner ring (IR) located between them. Each ring generally consists of 8 repeating subunits. Phenylalanine-glycine (FG)-rich repeats present in multiple Nups emanate inward from these rings to form the NPC’s central channel, and interact with transport factors to enable nucleocytoplasmic transport (5, 11). Apart from the CR, IR, and NR, the NPC features another prominent module known as the nuclear basket, also referred to as the basket (12–18). The basket is believed to play roles in mRNA transport and chromatin organization (14, 18–24). In yeast, Nup1, Nup2, Nup60, and Mlp1/Mlp2 are the main components of the basket and are referred to as basket Nups (4, 24–26). Their mammalian counterparts include Nup153 (ortholog of Nup60), Nup50 (ortholog of Nup2), and Tpr (ortholog of Mlp1/2) (12, 13). The basket has been observed in electron microscopy (EM) and in cryo-electron tomography (cryo-ET) studies of biochemically isolated NEs and NPCs of many organisms (15– 17, 24, 27), although the exact role of different Nups in the observed basket structures is not well defined. In these studies, the basket is described as an assembly consisting of eight struts emanating from the nuclear side of the NPC core that converge into distal densities, that could restructure and dilate to allow passage of large cargoes through the NPC (20). These studies also highlighted the need for further studies, including obtaining a 3D map of the basket in-cell and describing the molecular organization of basket Nups, along with exploring the structural dynamics of the basket and its neighboring peripheral NPC structures in-cell.

While the major observable features of the NPC, chiefly the basket and rings, have been known for decades in vertebrates, their organization and variability - from within a single cell to between species - has been largely undefined. However, recent work has highlighted that the NPC’s architecture may vary significantly both within and between species (5, 28). To explore the nature of such variations, we performed in-cell cryo-ET on NPCs of cells from three different and evolutionarily divergent eukaryotes: fungi (*S. cerevisiae*), mammals (*M. musculus*; mouse’ NIH3T3 cells), and parasitic protozoa (*T. gondii*)(Fig. S1A). The cells of these organisms were rendered amenable for in-cell cryo-ET through cryo-focused ion beam (FIB) milling (29–31), that produces lamellae thin enough for cryo-ET from vitrified cells (32). Our in-cell cryo-ET dataset represents one of the largest of its kind, comprising 1604 tilt-series. Through subtomogram analysis and 3D classification, we classified different variants of

NPCs across these organisms, yielding maps of one mammalian NPC (mNPC), one parasitic NPC (pNPC), and two distinct yeast NPC (yNPC) variants in-cell. The prefixes ‘m,’ ‘p,’ and ‘y’ have been used to denote mammals (mouse; M.musculus), and parasitic protozoa (*T. gondii*), and yeast (*S. cerevisiae*), respectively. From these maps, we discerned the basket architecture in the mNPC and one yNPC variant, shedding light on how the basket is organized on the NPC. Subsequently, we performed integrative modeling, incorporating a vast array of biochemical data on basket-Nups along with our in-cell maps, to model the molecular architecture of the basket-Nups. These maps and models provided us with a structural blueprint of how the basket forms and functions within the NPC.

## Results

### A stable basket is associated with a double NR

Our previous study revealed at least two populations of yNPCs within each cell: a major population with a single nuclear NR, and a lesser population carrying a double NR (5). We set out to further investigate the structural variants of NPCs in yeast in-cell using cryo-ET (Fig. S1A and methods). Through 3D classification of a large data set of in-cell NPCs, we found that ∼73% of NPCs in yeast during its log-phase growth possess a single NR, while the remaining have a double NR, which is consistent with the proportions estimated from quantitative fluorescence imaging (Fig. 1A-C, Fig. S1B, C) (5).

**Fig. 1.**
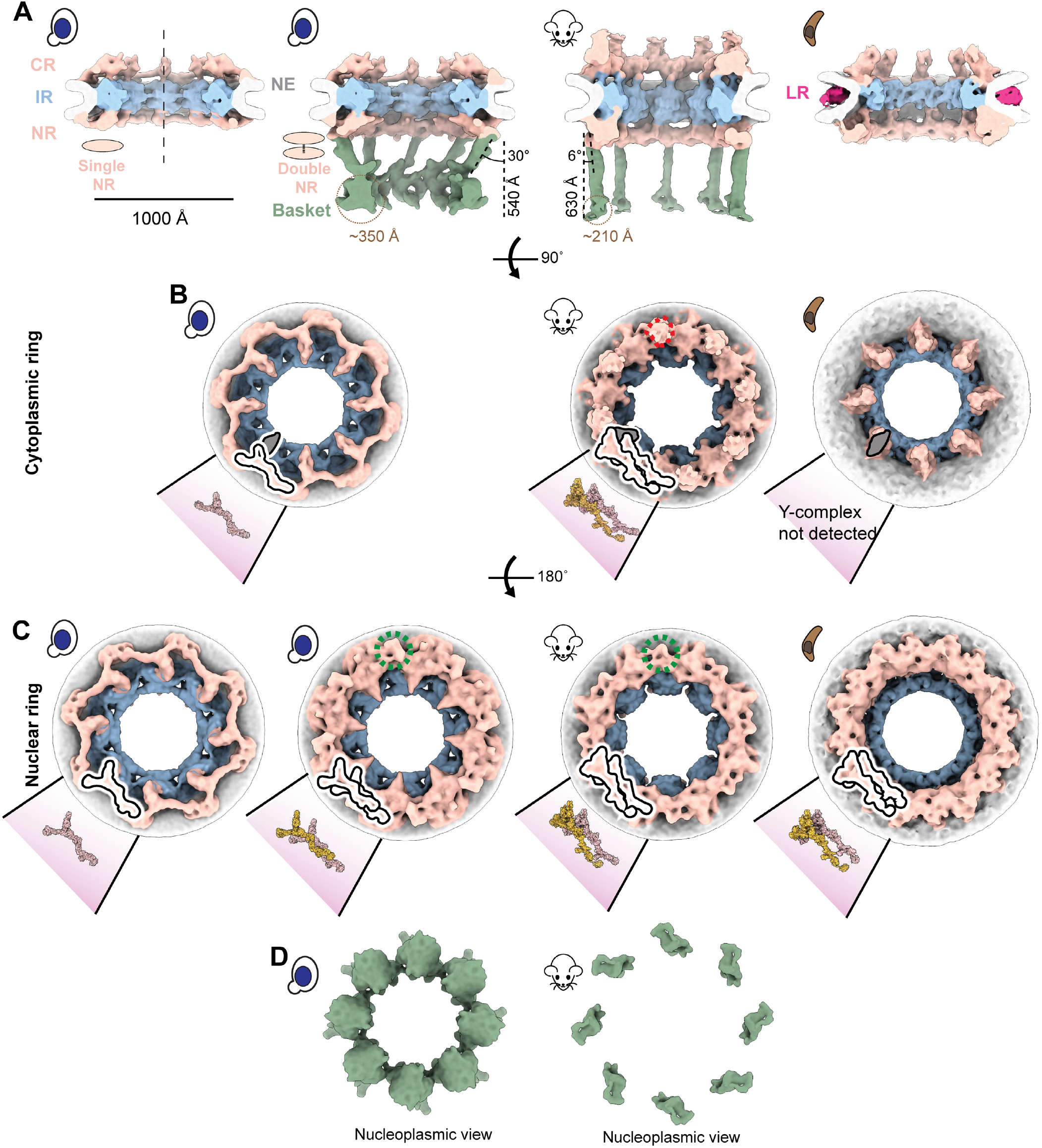
A stable nuclear basket is bound to a double nuclear ring. (**A**) Cross-sectional views along the central axis (dashed line) of the in-cell cryo-ET maps of yNPC with single and double NR variants, mNPC and pNPC. The nuclear basket is resolved for the mNPC and yNPC with a double NR. (**B**) Cytoplasmic views of the CR of yNPC with a single CR, mNPC with a double CR and pNPC with an incomplete CR. Also depicted are models of the single or double Y complex and the adjoining mRNA export platform, whose eight copies are arranged in head to tail orientation around the central axis to form the CR. Shown in dashed red lines is the region from which the cytoplasmic filaments emanate. (**C**) Nucleoplasmic view of the single and double NR of yNPC and double NR of mNPC and pNPC along with the models of the single or double Y complex. Shown in dashed green lines is the region in double NR from which the basket’s struts emanate. The models for yeast and mammalian CR/NR are from the PDB IDs: yNPC: 7N9F. mNPC/yNPC: 7R5J. (**D**) Nucleoplasmic views of the yeast and mammalian basket. CR: Cytoplasmic ring, IR: Inner ring, NR: Nuclear ring, NE: Nuclear envelope. LR: Lumenal ring.

The CR consists of eight subunits, each comprising one or two Y-complexes arranged in a head-to-tail orientation around the central axis passing through the center of the NPC (5, 6, 8, 33–35) (Fig. 1B, C). In addition to Y-complexes, the CR also has an mRNA export platform (5, 6, 8, 35–37) (Fig. 1B). The single NR also consists of eight Y-complexes, while the double NR carries sixteen such complexes in its two rings; the Y-complex rings proximal to and distal from the IR are referred to as the proximal and distal NR, respectively. The classification of the yNPCs into the single and double NR variants and their subsequent refinements gave us a more homogenous map of the single NR (devoid of any double NR densities). To date, a single NR is only observed in yNPCs (Fig. 1A, C). The IR and CR of the single and double NR variants are similar in stoichiometry and physical dimensions (Fig. S1B). Notably, a basket was observed only in the double NR variant in a dataset of more than 5100 NPC particles across 1449 tomograms from yeast, prompting the question of whether the single NR could support a stable and stoichiometric basket (Fig. 1A). However, it has previously been shown that most yNPCs, including both single and double NR forms, co-localize with basket components, with the exception of a class found only adjacent to the nucleolus that lack Mlp1/2 (23, 24), and that basket components in yeast are generally dynamic (5, 24, 38–40). Thus, we infer that basket components are more dynamic and so less well resolved in single NR NPCs (5, 38). Given that the basket was only resolved in the double NR variant in yeast we wondered if the mNPC, which always has a double NR (35) also has a discernible basket emanating from the double NR in a pattern similar to yNPC. Indeed, we also determined the architecture of the basket in mNPC and found it emanating similarly from its double NR (Fig. 1A, C). In an analogous manner to the basket and double NR, mNPC cytoplasmic mRNA export platforms emanate from the double CR, with an additional density likely representing the metazoan-specific Nup358 arrangement (41–43) (Fig. 1B).

### A protozoan NPC has a double NR and an incomplete CR

After observing the varying stoichiometries of the outer rings, we explored the diversity of these stoichiometries by performing in-cell cryo-ET and subtomogram analysis on NPCs of the protozoan parasite *T. gondii*, which diverged at least 1.5 billion years ago (44) and is responsible for toxoplasmosis in humans (45). Surprisingly, this protozoan NPC (pNPC) features a double NR but has a minimal CR which is discontinuous, even more so than the disjointed CR observed in the fungi *S. pombe* (Fig. 1A-C, Fig. S2) (46). Focused refinement of the pNPC’s subunits revealed that the incomplete CR appears to lack a full-length Y-complex, consisting only of a complex resembling the mRNA export platform (and perhaps under which is a remnant core of the Y-complex) (Fig. 1B). To date, no biochemical data exists to further rationalize the architecture of the CR of pNPC. Notably, pNPC has the smallest diameter observed so far under normal, i.e., non-stress and in-cell conditions. Its lumenal ring (LR) is more prominent, unlike the LR of yNPC and mNPC which are closer to the NE and thus more difficult to resolve (Fig. 1A). The LR of the NPC separates from the NE upon the NPC’s contraction and becomes distinctly more visible, as was also observed for contracted isolated yNPCs and yNPCs in-cell under cellular stress (5, 46).

### The basket consists of struts emanating from a double NR that end in a distal globular density

The basket consists of eight struts, each ∼100 Å thick, emanating from each of the eight subunits of the double NR at an angle from the central axis (30° for yBasket and 6° for mBasket), and terminating in a globular density referred to as the basket distal density (or basket ring), which is ∼540 Å and ∼630 Å away from the double NR in the yBasket and mBasket, respectively (Fig. 1A). The size of a single (one out of eight) basket distal density is ∼350 Å for the yBasket and ∼210 Å for the mBasket (Fig. 1A). In the yBasket, these densities are connected to form a ring with a diameter of ∼760 Å. However, no connections between distal densities were resolved in the mBasket with an apparent distal diameter of ∼1000 Å (Fig. 1A, D). These features of the basket bear broad similarities to those identified in isolated NEs and nuclei from various species (15, 16, 24, 27), except that the basket distal densities in isolated samples were mostly found to be connected, with their ring contracted unlike those in our in-cell maps (Fig. 1A, D) (15, 16, 24, 27).

### The nuclear periphery exhibits an exclusion zone around the basket

We leveraged the advantage of in-cell cryo-ET to examine the molecular environment in the proximity of NPCs with single and double NRs to explore why *S. cerevisiae* has two structural variants in NPCs and whether these variants had some specialized spatial distribution in the nucleus (Fig. S3A, B) (32, 47). In contrast to our expectation, in many cases, yNPCs with either a single or a double NR (those with a stable basket) were nevertheless found in similar environments, often adjacent to each other (Fig. S3A, B). One proposed role of the basket is to help create an exclusion zone around the NPC, presumably to streamline nucleocytoplasmic transport (21, 24). This hypothesis arose from observations of fixed and stained specimens in 2D electron micrographs (21, 24). Here, we generated a 3D average of the nucleoplasmic densities around the NPC to more clearly observe this zone and contextualize it with the basket structure. We used mNPCs, as they have morphologically better-defined heterochromatin than yeast (Fig. S3C). We generated a map of the mNPC, with a much larger box size, encompassing a significant portion of its surroundings, including nucleoplasmic regions (Fig. 2A). The map indeed revealed an exclusion zone around the mNPC and its basket on the nucleoplasmic side. This zone could also be seen directly in individual tomograms of mNPCs, which provide a clear depiction of this well-defined exclusion zone, which extends about 20 nm from the mBasket (Fig. 2A, B). Beyond the exclusion zone on the nucleoplasmic side, the densities likely represent lamina and chromatin, supporting the role of the NPC in chromatin organization (48, 49).

**Fig. 2.**
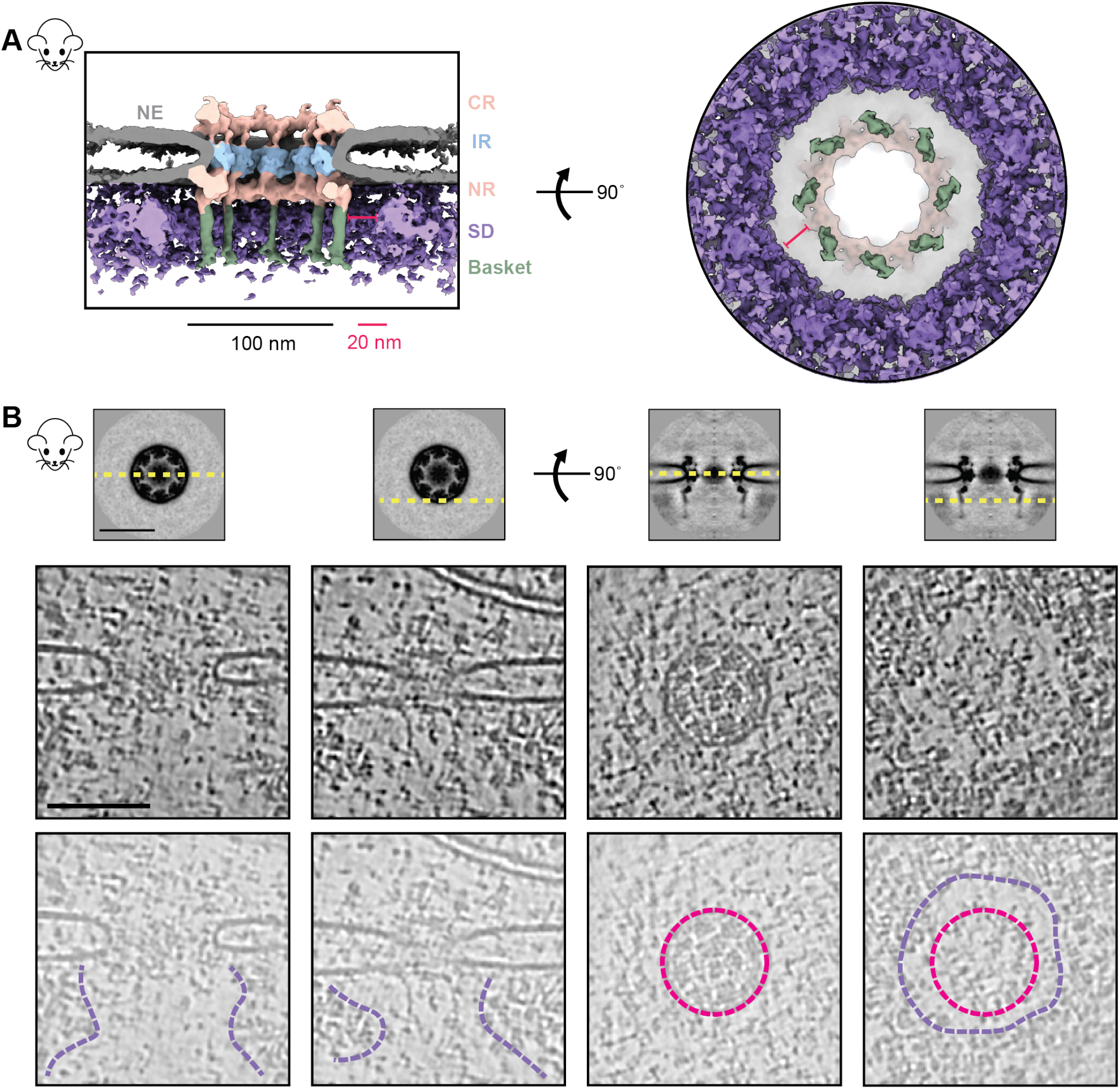
Direct observation of the heterochromatin exclusion zone around the mNPC. (**A**) The cross-sectional (left) and nucleoplasmic view (right) of the map of mNPC shows that molecular crowding (via surrounding densities; SD) around the mNPC is absent in the immediate vicinity of the basket (∼20nm).(**B**) Tomogram slices of different regions around the NPC show the extent of this exclusion zone. Top: slices of the mNPC average used to depict viewing planes (yellow, dashed) in tomogram slices below. Middle and bottom: tomogram slices of an mNPC viewed parallel (left) and perpendicular (right) to the plane of the NE, as depicted in the top panel. Tomogram slices are duplicated in the bottom row to show annotated views. Lumenal rings (pink) and boundaries of the exclusion zone (purple) are indicated. CR: Cytoplasmic ring, IR: Inner ring, NR: Nuclear ring, NE: Nuclear envelope. SD: Surrounding densities. Scale bars: 100 nm unless stated otherwise.

### The molecular architecture of the yeast and mammalian baskets

We used our iterative four-stage integrative approach to produce architectural maps of the yeast and mammalian baskets (Fig. 3 and Methods) (6, 50–54), based on subunit structure models, cryo-ET maps, chemical crosslinks, immuno-electron microscopy, coiled-coil propensities, sequence connectivity, excluded volume, and published data (Fig. 3 Stages 1,2) (SI Tables 1); a similar approach was used previously to determine the structure of the entire NPC (6). The modeling of the yBasket included basket Nups (yMlp, yNup1, yNup60, and yNup2, without their FG repeats) and NR Nups (yNup84 complexes), while the mammalian model consists of the basket Nups mTpr, mNup50, mNup153, and the NR mNup107 complexes (Fig. S4) (SI Tables 2, 3). The model optimizes the conformations and positions of these components while keeping the yNup84/mNup107 complexes fixed in their previously identified locations within our maps (Fig. 3 Stage 3) (SI Tables 2, 3). Before interpreting the models, we validated them using our standard assessment process (Fig. 3 Stage 4) (SI Tables 2, 3, Methods) (52, 54). Both models satisfy the data used to construct them (Fig. 3 Stage 1, Fig. S5, S6). In particular, the key input information, including the cryo-ET map, chemical croslinks, coiled-coil propensities, and subunit structure models is satisfied by a single cluster of structural solutions with an overall precision of 57 Å and 42 Å for yeast and mammalian, respectively (Fig. 3, Fig. S5, S6) (SI Tables 2, 3); the model precision is defined as the variability of the good-scoring solutions quantified by the average rootmean-square deviation (RMSD) of all solutions in the cluster. These precision estimates are considered when analyzing model features and comparing the two basket models.

**Fig. 3.**
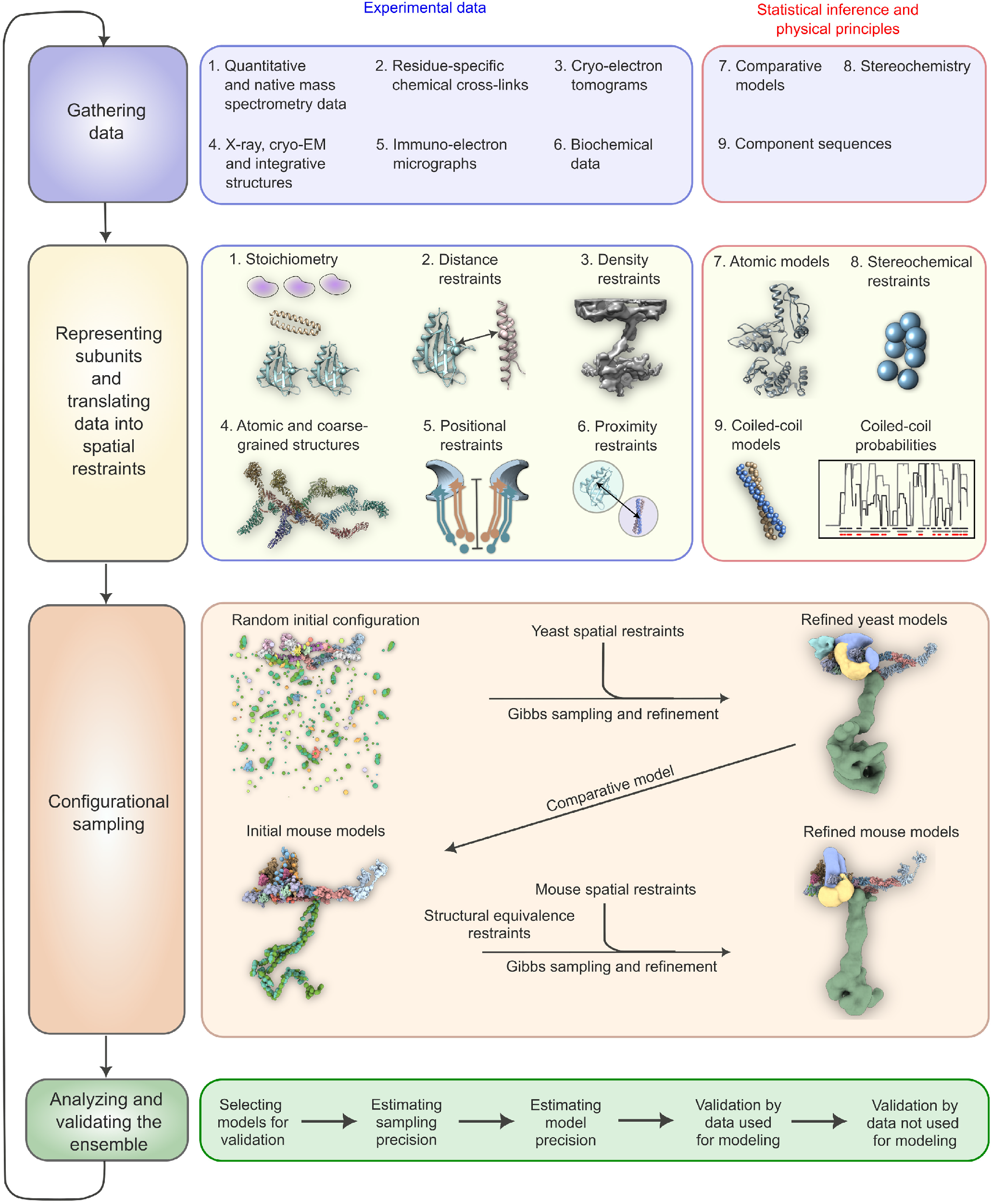
The four-stage scheme for integrative modeling of the baskets. Our integrative approach proceeds through four stages: (1) gathering data, (2) representing subunits and translating the data into spatial restraints, (3) configurational sampling to produce an ensemble of models that satisfies the restraints, and (4) analyzing and validating the ensemble. Stage 1 lists the experimental information used in this study for integrative modeling of baskets of yeast and mammals. Stage 2 lists representation and extracted spatial restraints obtained from information gathered in Stage 1. Stage 3 describes the configurational sampling to search for the models that satisfy the input information. A Gibbs sampling starting from a random initial configuration for the yBasket Nups generates the ensemble of good scoring models. The centroid model of the yBasket ensemble was used to model an initial mammalian basket. A similar Gibbs sampling with additional restraints from the mammalian data and structural equivalence restraint generates the ensemble of good scoring models. Stage 4 lists the model selection and validation protocol for the ensemble of good scoring configuration for both yeast and mammals.

### The yeast and mammalian basket models are similar in topology but different in the overall shape

The yeast and mammalian basket models have almost identical orthologous protein compositions (SI Tables 2, 3). The mammalian model was calculated to resemble the yeast model as much as possible while satisfying all the available mammalian data (SI Tables 2, 3, Methods). The two models share a similar topological arrangement, albeit with notable differences in the overall shape of the basket (Fig. 4A, C). To facilitate comparison, we dissected the basket into three modules, including the NR anchor (yNup1, yNup2, and yNup60 for yeast; mNup50 and mNup153 for mammals), basket strut (yMlp; mTpr), and basket distal modules (yMlp; mTpr) (Fig. 4B, D). The model revealed the proximity of the NR anchor module to the central channel of the NPC, NE, and NR (Fig. 4) and their association with the proximal NR via yNup60/mNup153 (Fig. 5A-D) (24). The N-terminus (blue) and C-terminus of yMlp/mTpr (red) are situated in the basket distal density, while the intervening region extends toward the double NR, forming the basket strut module. The yMlps/mTprs interact with the distal NR via yNup84/mNup107, consistent with the demonstrated requirement of yNup84 for anchoring Mlps onto NPCs (Fig. 5E, F) (24). The presence of two binding sites for the basket Nups, one on each proximal and distal NR, places the yNup60/mNup153 and yMlps/mTprs in direct interaction. This stabilizing interaction is feasible only in the context of the double NR, highlighting its importance in assembling a stable and less dynamic basket structure. However, our model also highlights how the basket might assemble in a single NR, as yMlps and yNup60 can interact with NR proteins (yNup84, yNup85, ySeh1) independently (Fig. 5A-D). Given the estimated model precision, our models are consistent with previously reported interactions between Nup60 and Nup2 (via their Nup60^N2BM^ domain) and Nup60 and Mlps (via their Nup60^MBM^ domain) (26, 55, 56). The model also revealed the presence of the coiled-coil domains of yMlp/mTpr in the basket strut module (Fig. 5E, F). The rod-like basket strut module exhibited around a 20° tilt between the yeast and mammalian baskets, resulting in a relatively large difference in the radius of the ring formed by the basket distal density (Fig. 4) (26). The basket distal module in both baskets is approximately globular, occupying the distal end of the basket (Fig. 4B, D). As is generally the case, it is unclear whether the differences between the yeast and mouse basket models reflect the differences between species, experimental conditions for purification and structure determination, and/or functional states.

**Fig. 4.**
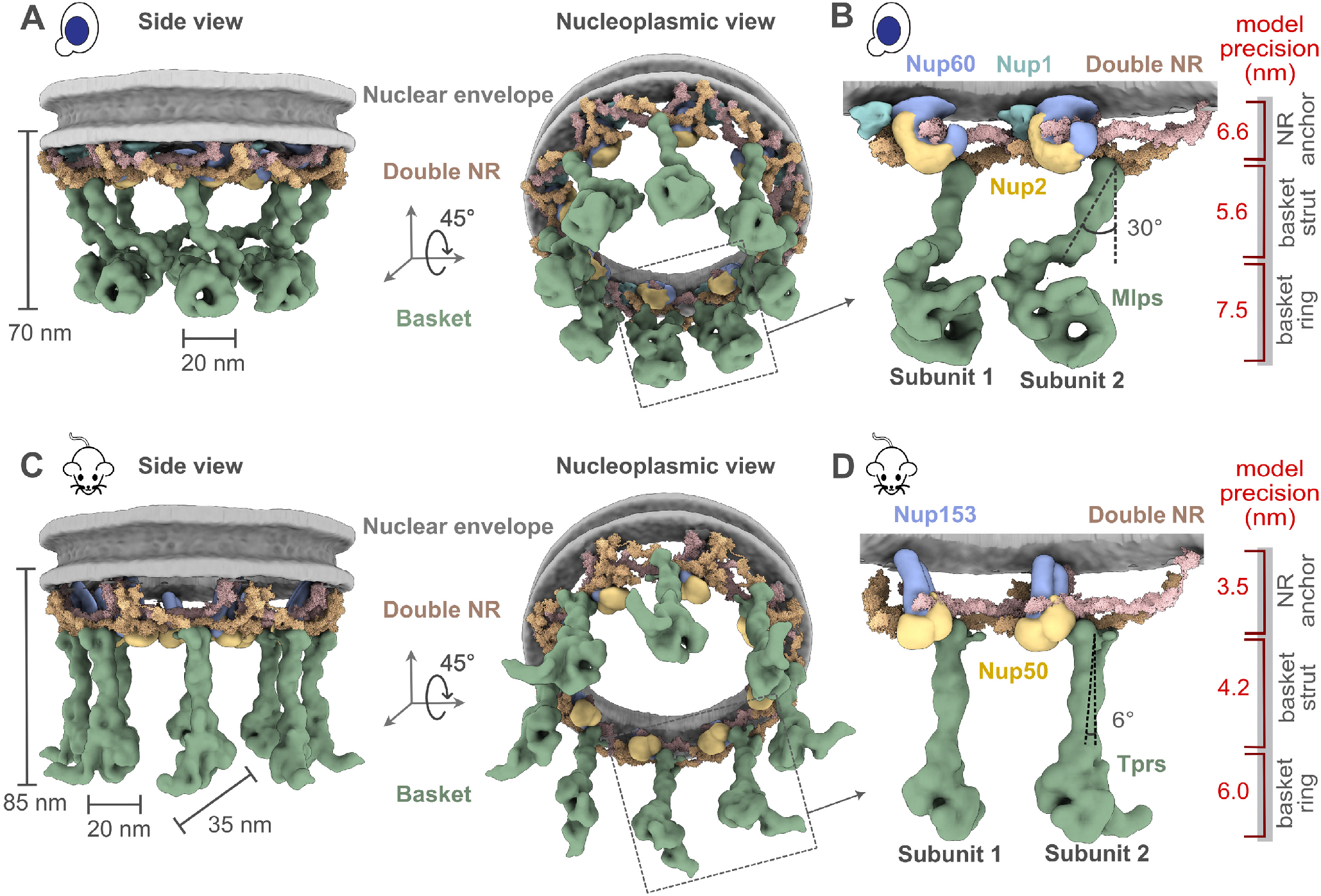
Integrative structure models and precisions of yeast and mammalian nuclear baskets. (**A, C**) Localization probability density for the yBasket (containing yMlp, yNup1, yNup60, yNup2 Nups) and mBasket (containing mTpr, mNup50, mNup153 Nups) obtained from the ensemble of good scoring models. A localization probability density map for a set of models is defined as the probability of observing a model component at any point in space. The yMlps/mTprs (green) are attached to the double NR and NE (light gray half-toroid) with a common interacting FG Nups yNup60/mNup153 (violet). (**B, D**) A close-up view of the two subunits of the yBasket and mBasket with model precision for different basket segments. Shown here are yMlp/mTpr (green), FG Nups yNup1 (cyan), yNup2/mNup50 (yellow), and yNup60/mNup153 (purple).

**Fig. 5.**
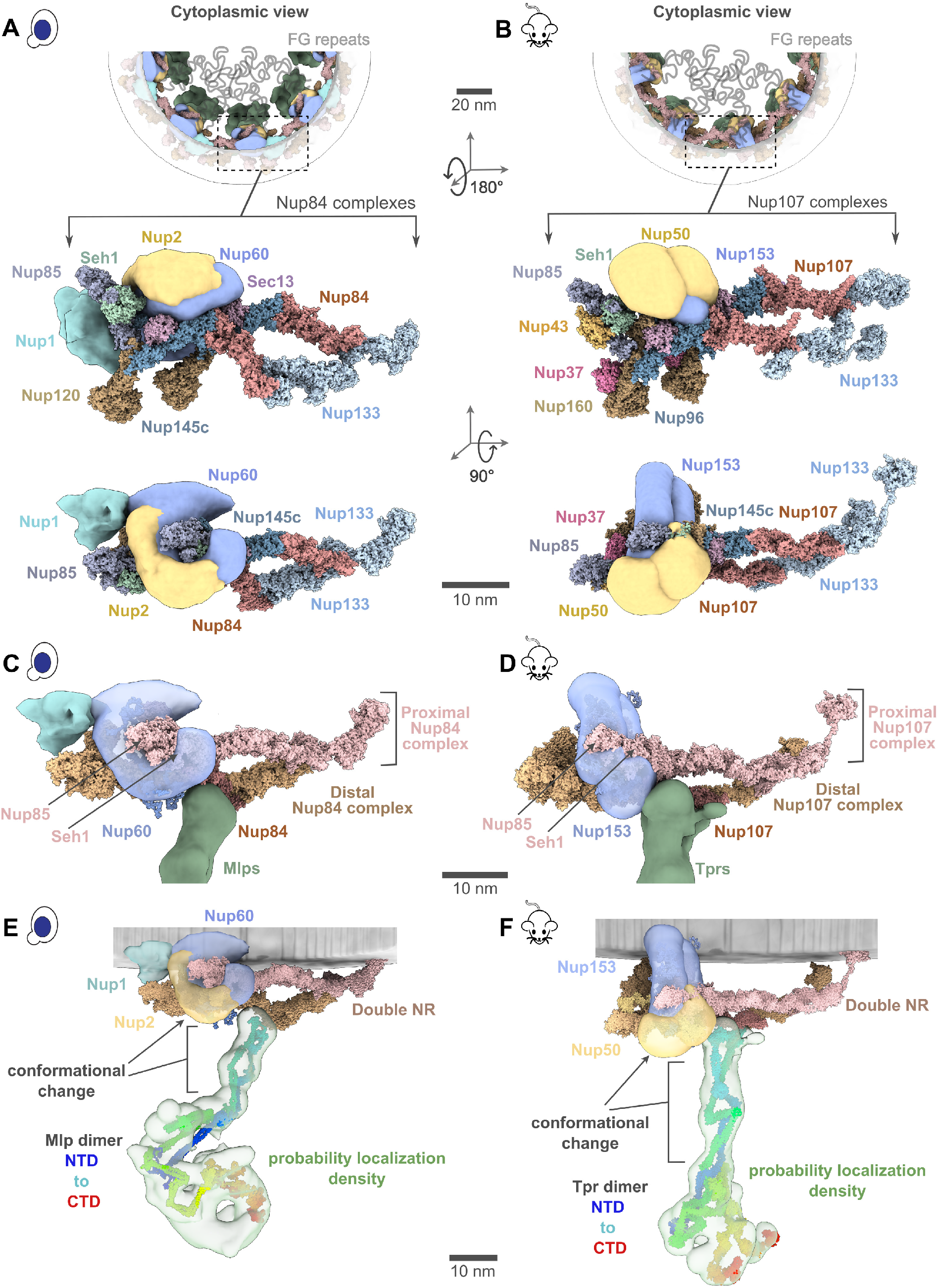
Position of different nucleoporins in the basket model. (**A, B**) Cytoplasmic view of the yeast and mammalian basket zooms into a Y-complexes (yNup84 complex for yeast and mNup107 complex for mammalian). Schematic representations of the FG repeats and anchoring positions are shown as curly lines (gray). Each double Y-complex’s zoomed image highlights individual Nups with two different views. Localization densities of the basket anchors, yNup1 (cyan), yNup2/mNup50 (dark yellow), and yNup60/mNup153 (violet), are shown relative to the Y-complex Nups. (**C, D**) Close-up inner view of the localization densities of the yNup60/mNup153 (violet) that contacts with the proximal nuclear ring (pink), whereas the localization densities of yMlp/mTpr dimer (green) contacts the distal nuclear ring (tan) of the NPC. yNup2/mNup50 not shown in this view. (**E, F**) The yMlp/mTpr dimer centroid models were colored from the N-terminus (blue) to the C-terminus (red) and shown embedded within their localization probability density (light green). The unidirectional arrows indicate local conformation change of yMlp/mTpr and yNup60/mNup153 between ybasket and mbasket models.

### The FG regions in the NR anchor module face the central channel

The yNup1 and yNup2 had not been included in the previous model of the NPC, due to the lack of data about their positions (5, 6, 50). In our models, the non-FG regions of FG Nups in the NR anchor module localize between the two copies of Nup85 in the proximal and distal yNup84/mNup107 complexes, with a precision of 6.6 nm (Fig. 4B, D and Fig. 5A, B). Both copies of yNup2 and yNup60 (and their mammalian orthologs) are proximal to each other as well as to several Nups in the double NR (Fig. 5A, B). The yNup1 is also positioned near yNup60, anchoring both to the nuclear envelope, consistent with previous mapping (55). This arrangement exposes the NR anchor module to the central channel. By virtue of the position of the globular anchor domain of the FG Nups, the FG anchoring sites are positioned such that the FG repeats face into the central channel, as is the case for previously localized FG repeat regions (5, 6, 57). This observation serves as further validation of our models, as this feature was not imposed on the models (Fig. 5A, B). The anchors are not sufficiently long to extend far into the struts or into the distal basket, indicating that the associated FG repeats remain localized to the proximal end of the basket and so spatially segregating transport from the initial docking processes occurring at the distal basket (58, 59).

### The resilience of the basket architecture may arise from a combination of rigid and flexible modules

It was proposed that the structural resilience of the NPC is achieved via an architecture that combines flexible and rigid modules, akin to the design of a suspension bridge (5, 6). Our models indicate that this architectural principle extends to the basket. Specifically, the apparently rigid components (SI Tables 2, 3) of the model include the subunits of the double NR (pink and tan, Fig. 5C, D), structured domains of the NR anchor module, and the basket strut module (green, Fig. 5C, D). The apparently flexible components (SI Tables 2, 3) of the model include the unstructured domains of the NR anchor module (violet, Fig. 5C, D) and the basket distal module (green, Fig. 4B, D). Our model indicates that the N-terminal amphipathic helices (Fig. S4C) within the yNup60/mNup153 of the NR anchor module may serve as critical anchors to the nuclear envelope (gray, Fig. 4) in full agreement with previous findings (26, 55). The flexible regions of the NR anchor module connect to the rigid double NR and basket strut module, mimicking the role of suspension cables that connect rigid columns and the roadway of a suspension bridge (55). The suspension bridge-like architecture may provide the necessary resilience of the basket while transporting large cargos.

## Discussion

The array of stoichiometries we observe for the outer rings (CR+NR) across and within species, ranging from incomplete to 1 to 2, highlights the NPC’s modular construction and its structural plasticity, which allows it to easily adapt to gain or lose additional subcomplexes, presumably to confer alternate functionalities. *T. gondii*’s NPC appears to have the full mRNA export platform but an incomplete CR, suggesting that the full Y complex in the CR might be more dispensable than the mRNA export platform. While *T. gondii*’s CR is quite distinct, its IR is similar to other IRs, consistent with the differences among NPCs of various organisms being more pronounced in their outer rings rather than the IR, which appears to represent the most structurally and evolutionarily conserved module of the NPC (60). The NPCs with alternative stoichiometries of rings can be used to understand how the NPC’s parts are formed (e.g., we show here how the distal NR could help bind and stabilize the basket) and their dispensability (e.g., an NPC can function without a full length Y-complex on its CR, as shown here for *T. gondii* and has been shown for *S. pombe* (46)). Like the CR, the NR can also show different copy numbers. However, in the organisms examined so far, there is at least one complete NR as a necessary minimum for NPCs. Beyond this, many organisms can carry two NRs on either some or all of their NPCs; and Dictyostelium NPCs may usually carry 3 such NRs (61). The reason for this variability in NR copy number is not immediately evident; however, functional insights may be gained by examining the relationship between the NR architecture and that of the nuclear basket.

It should be noted that the term *‘basket’*, as traditionally defined and understood, refers to a stable architecture with struts. However, as we have shown and is consistent with observations here, the basket can be highly dynamic, and can also interact with many other proteins and indeed may have additional components, such as yPml39 / mZC3HC1 (56, 62). Various studies have established that much of the pool of nuclear basket components is dynamically associated with the yNPC (38); and, similar to yeast Nup1, Nup60, and Nup2 (39, 40), mammalian Nup153 and Nup50 are also dynamically associated with their respective NPCs (63). However, the major strut component in yeast (Mlp1/2) is dynamically associated (39, 40), whereas that in mammals (Tpr) seems more stably associated with the NPC, in agreement with the near ubiquitous appearance of the basket in mNPCs (above) (12, 13, 64). Moreover, we found that every double NR yNPC had a clearly associated basket, whereas the single NR NPCs did not have clear morphologically discernable baskets. However, in seeming contradiction, most yNPCs associate with basket components *in vivo* (6, 23–25, 38, 65). A reconciliation of this apparent contradiction can be made simply, as follows. We know that the majority of the yeast Mlp1/2 pool is also dynamic (above); we therefore suggest that for the single NR yNPCs, their association with basket components, including Mlp1/2, is transient, and furthermore that the basket components are more flexible and perhaps not stoichiometric (17), such that their presence is difficult to establish by in-cell cryo-EM. In contrast, we suggest that the double NR yNPCs, by virtue of their possession of an additional set of basket protein binding sites in their extra NR, can bind much more stably to Mlp1/2, thus accounting for the ubiquitous observation of basket struts in the double NR yNPCs. This idea also agrees with the observation, using scanning electron microscopy, of complete baskets on only a subset of yNPCs, although all had some strut-like nuclear filaments (17). We may also propose a function as to why yeast possess these minority double NR yNPC forms; while most yNPCs (with single NRs) utilize a rapidly reversible recruitment mechanism for the basket during mRNA export (38, 65), as masters of *‘bet hedging’* (66) yeast keep in reserve an NPC subset with preassembled NPCs, perhaps to accommodate rapid changes in mRNA export or to ensure maintenance of epigenetic memory (67). In mNPCs, the presence of a double NR on essentially all NPCs ensures the stable association of Tpr and a morphologically recognizable basket.

When an NPC acquires a double NR, it not only becomes more stably associated with Mlp1/2; it also apparently becomes capable of more stably binding yNup1, yNup60 and yNup2, as the presence of the extra ring provides observed additional binding sites for these proteins on the NPC (Fig. 5A). The binding of these proteins to this aforementioned hub in the NE has been shown to be managed by posttranslational modifications (68, 69). Interestingly, a similar phosphorylation-driven mode of assembly and disassembly is found in flexible connector-containing Nups of cells that undergo NE breakdown, as we previously suggested (5).

Our cryo-ET analysis was not able to resolve a basket in the double NR-containing *T. gondii* NPCs, most likely due to the limited data set, and thus we cannot at this stage rule out the presence of a highly divergent basket structure or loss of the basket, as we were unable to identify obvious Tpr homologs in this organism. However, this may also suggest that the presence of a double NR might not be sufficient to sustain a highly stable basket assembly. Indeed, even in yeast and vertebrates the presence of a double NR does not guarantee the continued presence of a full stable basket, as it has been shown that under stress (70, 71) or under certain cellular states (68, 69, 72), the nuclear basket can dissociate from the NPC.

It was previously observed that the C-terminus of Mlp1 acts as a necessary transient docking site for messenger ribonucleoproteins (mRNPs) during mRNA export (59) and that the nucleocytoplasmic transport of large molecules through the NPC involves an increase in the basket ring radius (20). Our model rationalizes these observations as follows. The C-terminus of Mlps is located in the basket distal density. If the difference between the yeast and mammalian models was indicative of the basket structural dynamics, the dynamics of the basket would involve the movements of the basket subunit and conformational changes of anchor Nups (Fig. 5E, F) (24, 73). Moreover, the flexible linkers connecting the coiled-coil segments from the yMlps/mTprs dimer in the struts may also afford the flexibility to the basket to contract and expand as seen for other rings (5, 46). Such motions could account for the expansion and contraction of the basket seen during passage of large cargoes (20). A significant portion of the basket distal density in yeast remains unaccounted for in our integrative model, possibly indicating the presence of cargo, transport factors or other elements, which aligns with the observation that this density could be a docking site for cargo (Fig. 5E, F). Unlike the yNPC, the basket distal density in the mNPC is considerably smaller, and thus, a larger proportion of its density is accounted for in its models (Fig. 5E, F).

The sizes of many mRNPs range from ∼200 to 600 Å, smaller than the diameters of the rings, including that of the basket, potentially allowing them to pass unaltered across the NPC (74, 75). However, some mRNPs can have elongated shapes or be much larger than the basket’s diameter (20, 74, 75); in these cases, their shape can be adjusted or remodeled to pass through the NPC, and the basket too can remodel as seen during passage of such large cargoes (20, 76). These adjustments or remodelings could commence while the mRNP is docked at the basket distal density. The role of the basket has primarily been discussed for nuclear export, not import. However, it would be interesting to explore what happens to the basket during import, including for large cargoes like HIV capsids that can pass through the NPC as intact entities (77).

Apart from chromatin exclusion, NPCs also influence chromatin organization through the context-specific localization of either active or repressed genes to the NPC (78–81). These localizations require basket Nups such as yNup1, yNup2, yNup60, mNup153, and Mlps, implying the basket’s critical role in these localizations (78–81). Similar to mRNA docking, these localizations may also occur at the basket distal density, consistent with reported interactions between the C-termini of Mlps (which we show to form the distal density) and complexes involved in genes localization to the NPC (82). Interestingly, chromatin organization is not only impacted by the presence of the basket but also by its systematic absence, as shown recently that basketless yNPCs are involved in a process of subtelomeric gene silencing (83).

## Acknowledgments

We thank the members of the Villa, Sali, and Rout labs, as well as Chris Akey and Steve Ludke for their feedback and support. D.S. was supported by a Damon Runyon Postdoctoral Fellowship (DRG-2364-19) and is currently supported by the NIH’s K99 Pathway to Independence Award (K99AG080112). J.H. was supported by an EMBO longterm postdoctoral Fellowship (ALTF 871-2020). E.V. is an investigator of the Howard Hughes Medical Institute. We acknowledge the use of the UC San Diego cryo-EM facility, which was built and equipped with funds from UC San Diego and an initial gift from the Agouron Institute. This work was supported by an NSF MRI DBI 1920374 (to E.V.), U54 AI170856 (to E.V.), NIH P41 GM109824 (to M.P.R., B.T.C. and A.S.), NIH R01 GM083960 (to A.S.), NSF-1818129 and Spanish Ministerio de Ciencia e Innovacion PID2020-116404GB-I00 (to J.F.-M.), NIH R01 GM112108 and NIH GM117212 (to M.P.R.). We thank Prof. Ricardo Henriques’s Lab for LAT_E_X template used for typesetting this document.

## Author Contributions

- <u>Conceptualization:</u> D.S., N.S., J.H., M.P.R., A.S., and E.V.
- <u>Sample Preparation and Data Acquisition:</u> D.S., J.H., F.S., Z.L., S.S., J.F-M., T.v.E., K.M, Y.S., J.W.
- <u>Analysis:</u> D.S., N.S., J.H., I.E., M.D., F.S., Z.L., Q.G., J.F-M., K.M., B.C.
- <u>Writing:</u> D.S., N.S., J.H., I.E., J.F-M., M.P.R., A.S., and E.V. with input from all the authors.

## Methods

### Cell culture, vitrification and sample preparation

W303 yeast cells were cultured in yeast extract peptone dextrose (YPD) media supplemented with adenine hemisulfate. These cells in the log-growth phase were collected and deposited on glow-discharged Quantifoil grids (R 2/1, Cu 200-mesh grid, Electron Microscopy Sciences), as described previously (5). The mouse fibroblasts cells (NIH3T3) were cultured at 37°C and 5% CO2 in DMEM with 10% fetal calf serum. Cells were seeded onto glow-discharged and Fibronectin-coated Quantifoil grids (R1/4, Au 200-mesh grid, Electron Microscopy Sciences). Following this seeding, the cells were cultured for 2 more hours on the grids to allow for their stable adherence onto the grid. In some cases, grids were micropatterned with 40 μm circles and treated with 100 nM jasplakinolide for a further two hours after seeding. The tachyzoites (*T. gondii* in rapid growth phase) were thawed out from liquid nitrogen and cultivated in human foreskin fibroblasts (HFFs) using Dulbecco’s modified Eagle’s medium (DMEM), with medium changes every 12 to 24 hours. To collect tachyzoites, trypsin-treated, parasite-infected HFFs were mechanically disrupted using a 27-gauge syringe, and the mixture was filtered to separate tachyzoites from HFF debris. The tachyzoites were then centrifuged, resuspended in DMEM with 30% FBS and 10% DMSO, and deposited on EM grids for vitrification, as described previously (84). Excess media was manually blotted from the back (opposite to the carbon film and seeded cells). Grids were plunge-frozen in a liquid ethane-propane mixture (50/50 volume, Airgas) using a custom-built vitrification device (Max Planck Institute for Biochemistry, Munich). Frozen grids were clipped into AutoGrids with a milling slot (Thermo Fisher Scientific) to allow milling at shallow grazing angles as described previously (31, 85). Cryo-FIB milling was performed in an Aquilos Dual-Beam (Thermo Fisher Scientific) as described previously (31, 85).

### Tilt series acquisition

Tilt series were acquired on the Titan Krios G3 (Thermo Fisher Scientific) at 300 keV with either a K2 detector and Quantum 968 LS post-column energy filter or a K3 Summit detector with 1067HD BioContinuum post-column energy filter in counting and dose fractionation modes (Gatan). The tilt-series parameters were as follows: tilt range: ± 45-60°, pixel size of 3.45 Å (yeast), 1.32 Å (mouse fibroblasts), 3.328 Å (*T. gondii*), tilt increment: 3° (higher for some samples), effective defocus range: -2 to - 11 μm, total fluence: ∼100-180 e-/Å2. All image acquisition was done using SerialEM software (86, 87). For some tilt-series, parallel cryo-electron tomography (PACE-tomo) scripts were used (88). In total, 1449, 136, and 19 (total: 1604) tilt series were used for yeast, mouse, and *T. gondii*, respectively. This data set included 153 tilt-series of yeast from EMPIAR-10466 (8).

### Subtomogram analysis

Frames of the tilt images were motion-corrected using whole-frame motion and organized into stacks in WARP (89, 90). The motion-corrected tilt series were then aligned in AreTomo (91). The aligned tiltseries stacks were subsequently re-imported into WARP for CTF estimation, defocus handedness determination, and final reconstruction (89, 90). The CTF estimation and defocus handedness were manually inspected and further refined as needed. In tomograms, nuclear pores were manually picked in IMOD (92). For each pore, in addition to the coordinate of the pore’s center, an additional point approximately 50-100 nm on the cytoplasmic side was marked. The pores were oriented using these two points with the Dynamo dipole picking mode (93). The subtomograms of the pores, with these initial orientations, were generated in WARP at a pixel size of 10 Å. The total number of pores picked were ∼5160 for yeast, ∼220 for mouse, and ∼50 for *T. gondii*, respectively. A small number of pore particles were used to generate a C8 symmetrized initial model in Relion (94). This initial model served as a reference for refining all the pore particles with C8 symmetry. The refinements were performed with local searches around the initial orientation (initial Euler angles), using the sigma_ang/rot/tilt/psi parameters to restrict the angular searches and prevent the pore particles from flipping. The term sigma_ang/rot/tilt/psi in Relion specifies the width of the Gaussian prior on the starting Euler angles. 3D classification (without alignments, simply referred to as classification), using C8 symmetry, was performed using a mask focused on the inner ring of the NPC to select good particles and discard bad ones. The selected NPC particles were refined further with C8 symmetry. For yeast, classification was performed using a mask focused on the nuclear ring to classify out NPCs with single and double NR, which accounted for ∼77% and the remaining ∼23% of total NPCs, respectively. The symmetry expansion was carried out to isolate subunits of the NPCs. These subtomograms of the subunits were then reconstructed at a pixel size of 10 Å in WARP. The relion_reconstruct was used to generate an average of these subtomograms for use as reference in the refinement of these subunits using a mask focused on the IR subunit. Following refinement, classification was performed, using the mask focused on the IR subunit, to select good subunits and discard bad ones. The refinements of the good subunits of IR, CR, NR, and the basket (as applicable) were then performed using their respective shape masks. All the refinements and classification of subunits were done without the use of symmetry. The total number of subunits used in the final refinements were ∼28600 for yeast (out of which,∼6600 were from the NPC with double NR), ∼800 for mouse, and ∼265 for *T. gondii*, respectively. The 0.143-cut-off criterion of the Fourier Shell correlations (FSC) between masked and independently refined half-maps was used to estimate all the reported resolutions (95). The maps of the subunits of these different rings were composited to generate the final map of the entire subunit of the NPC. This composite map was fit into the map of the whole NPC (of C8 symmetry) using Chimera’s fit-to-map tool (96), and then C8 symmetrized using relion_image_handler. The entire processing of the data from separate organisms was done completely separately and independently. The schematic of the entire workflow and resolution estimates is also shown in Figure S1. v3.1.1 of Relion was used for all steps involving Relion (94). v1.09 or v1.1.0-beta1 of WARP was used for all steps involving WARP (89, 90).

### Pairwise distances amongst yNPCs and their radial distribution function [g(r)]

The coordinates of yNPCs with single or double NR in their tomograms were obtained following their subtomogram analysis. For each tomogram, pairwise distances among all yNPCs, as well as those with single and double NR, were calculated using these coordinates. These distances were then used to estimate the g(r) for each tomogram. The g(r) values from all the tomograms were averaged to generate the final g(r) shown in (Fig. S2B). It should be noted that these pairwise distances and their corresponding g(r) values are averages for all yNPCs and might not apply to small subsets of yNPCs. For instance, yNPCs near the nucleolus are likely to be less enriched in double NRs (with a stable basket). This observation comes from fluorescence imaging, which has shown that yNPCs near the nucleolus lack yMlps (one of the basket-Nups) and have a low level of NR-Nups, indicating a preference for single NR without the basket (5, 23, 38, 97).

### Chemical cross-linking and MS (CX-MS) analysis of affinity-purified yeast NPCs

CX-MS of Mlp1-PPX-PrA tagged, affinity purified, native, whole NPCs have been described in detail in (5, 6). To expand and complement these datasets with cross-links mapping exclusively to basket Nups fully assembled into the NPC, we used NPCs affinity purified using Dbp5-PPX-GFP and Gle1-PPX-PrA as the handles using a similar protocol, with the following modifications: After native elution, 1.0 mM disuccinimidyl suberate (DSS) was added and the sample was incubated at 25°C for 40 minutes with shaking (1,200 rpm). The reaction was quenched by adding a final concentration of 50 mM freshly prepared ammonium bicarbonate and incubating for 20 minutes with shaking (1,200 rpm) at 25°C. Crosslinked NPCs were pelleted by spinning for 20 minutes in a TLA-55 rotor (Beckman) at 25,000 rpm. The pelleted samples (∼50 mg) were resuspended in 1xLDS with 25 mM DTT and incubated at 70°C for 10 minutes. Reduced samples were alkylated by adding a final concentration of 100 mM iodoacetamide and incubating in the dark at 25°C for 30 minutes, followed by the addition of an additional 25 mM DTT and further incubation for 15 minutes. Alkylated and reduced samples were denatured at 98°C for 10 minutes and then loaded into 4% SDS-PAGE Bis-Tris gel and run for 10 minutes at a constant 120 V to reduce the complexity of the sample. For in-gel digestion, the high-molecular-weight-region gel bands corresponding to cross-linked NPC proteins were sliced and proteolyzed by trypsin as previously described (6). In brief, gel plugs were crushed into small pieces and 5–10 μg of sequencing-grade trypsin (Promega) per ∼100 μg protein were added. Trypsin was supplied in two equal additions and incubated with gel pieces at 37°C in 50 mM ammonium bicarbonate, 0.1% (w/v) Rapigest (Waters). After the first addition, the samples were incubated for 4 hours. After the second addition, the samples were incubated overnight. Peptides were extracted by formic acid and acetonitrile, and dried partially by vacuum centrifugation. To remove the hydrolytic insoluble by-products of Rapigest, the sample was centrifuged at 20,000 g for 10 min. The solution was transferred to another tube and then further dried by vacuum centrifugation. Peptides were separated into 6-7 fractions by high pH reverse phase fractionation in a pipet tip self-packed with C18 resin (ReproSil-Pur 120 AQ, 3μm, Dr. Maisch GmbH). Each peptide fraction was resuspended in 5% (v/v) methanol, 0.2% (v/v) formic acid and loaded onto an EASY-Spray column (Thermo Fisher Scientific, ES800, 15cm x 75mm ID, PepMap C18, 3mm) via an EASY-nLC 1200 (Thermo Fisher Scientific). The column temperature was set to 35°C. Using a flow rate of 300 nl/min, peptides were gradient-eluted (3–6% B, 0-6 min; 6–34% B, 6-97 min), where mobile phase B was 0.1% (v/v) formic acid, 95% (v/v) acetonitrile and mobile phase A was 0.1% (v/v) formic acid in water. An Orbitrap Fusion Lumos Tribrid (Thermo Fisher Scientific) was used to perform online mass spectrometric analyses. Full MS scans were performed at least every 5 s. As time between full scans allowed, ions with charge states +4 to +8 were fragmented by higher-energy collisional dissociation in descending intensity order with a maximum injection time of 800 msec. Both precursors and fragments were detected in the Orbitrap. The raw data were searched with pLink2 (98) with cysteine carbamidomethyl as a fixed modification and methionine oxidation as a variable modification. The initial search results were obtained using a default 5% false discovery rate (FDR) expected by the target-decoy search strategy. Spectra corresponding to basket components were selected and manually verified to ensure data quality (6).

### Integrative modeling of the basket

Coarse-grained structural models of the yeast and mouse baskets were computed using an integrative modeling approach (6, 50–54), based on information from varied experiments, physical principles, statistical preferences, and prior models (SI Table 1). The yBasket model includes the yMlp1/2, FG Nups (yNup1, yNup2, and yNup60), as well as the double NR Nups (yNup120, yNup85, yNup145c, ySec13, ySeh1, yNup84, and yNup133) (4, 24–26). The mBasket model includes the orthologs of yeast Nups (mTpr, mNup50, mNup153, mNup160, mNup85, mNup96, mSec13, mSeh1, mNup107, mNup133, mNup43, and mNup37) (12, 13). Modeling positioned the yMlp/mTpr and FG Nups relative to the fixed double nuclear ring; in addition, it optimized the conformations of the disordered Nup regions. The modeling protocol was scripted using the Python Modeling Interface (PMI) package version a41075a, which is a library for modeling macromolecular complex structures based on our open-source Integrative Modeling Platform (IMP) package version 2.19 (https://integrativemodeling.org) (52).

### Stage 1: Gathering information

The sequences of the basket Nups were obtained from the Uniprot database (99) (SI Tables 1, 2). Their stoichiometry in the yNPC was previously determined by quantitative mass spectrometry of the isolated yNPC complex (6) (SI Table 2). In total, 626 unique intra- and intermolecular DSS cross-links were previously identified using mass spectrometry (6, 24, 100). The cryo-ET map described here informed the overall shape of the basket and its anchoring on the double nuclear ring. The structural model of yNup84 complex of the double NR was previously determined by an integrative approach (5). The structural model of the yMlps was informed by the coiled-coil propensities and heptad repeat alignments and was generated using COCONUT software (101) (Fig. 3 stage 2 and Fig. S4A, B) (SI Table 1). The yNup2 structural model was obtained from the AlphagFold database version 4 (SI Table 2). Direct physical interactions between yNup60, yNup2, and yMlp1 were determined by *in vitro* binding assays (55) (SI Table 1). Previously determined immuno-electron microscopy images help localize the terminal domains of the yMlps (24). Similar information was used for mBasket modeling (12, 102) (Fig. 3 and Fig. S4) (SI Tables 1, 3).

### Stage 2: Basket representation and spatial restraints. Basket representation

We modeled only a single subunit out of the eight subunits comprising the entire yBasket, mostly without explicitly considering the interfaces between the eight symmetry units in the yNPC. This simplification was possible because the explored positions and conformations of the basket components do not clash with each other across the symmetry unit interfaces, courtesy of their anchoring on the fixed double NR. The stoichiometries of yMlp1 and yMlp2 are ambiguous (6). Thus, we used two copies of poly-alanine per symmetry unit (yMlp), representing both yMlp1 and yMlp2; the yMlp length was set to that of yMlp1. The model also included a single copy of yNup1 and two copies of yNup2, yNup60, and the heptameric yNup84 complex. FG repeats were not included in the model. In total, the yBasket model consists of 21 protein subunits of 12 types (SI Table 2). A similar representation was used for mBasket modeling with two copies of mTpr, two copies of mNup50, mNup153, and the nonameric mNup107 complex (SI Tables 1, 3) (35, 103, 104). Thus, the mBasket model consists of 24 protein subunits of 12 types (SI Table 3). Each component was represented in a multiscale fashion to balance the accuracy of the formulation of restraints and the efficiency of structural sampling (Fig. S4C) (SI Tables 2, 3).

### Spatial restraints on yeast and mouse baskets

The subset of input information was converted to spatial The subset of input information was converted to spatial restraints for scoring alternative models (52). These restraints include upper bounds on pairs of crosslinked residues based on chemical crosslinks, the correlation coefficient between Gaussian Mixture Models of a model and the cryo-ET map, positional restraints on NTD/CTD domains of yMlps based on immuno-EM localizations, distance restraints between pairs of domains based on affinity co-purification data, positional restraints on residue segments predicted to lie within the nuclear envelope, connectivity restraints between consecutive pairs of beads in a subunit, excluded volume restraints between non-bonded pairs of beads (SI Table 2) (52). For the mBasket (SI Table 3), crosslinking and affinity copurification data were unavailable. However, we supplemented the remaining mBasket restraints with structural equivalence restraints; these distance restraints are designed to maximize the similarity between the mouse and yeast models across the aligned residues, subject to the satisfaction of the remaining restraints.

### Stage 3: Structural sampling

The initial positions and orientations of rigid bodies and flexible beads were randomized except for the double NR rigid body (SI Table 2), whose position was obtained by fitting into the cryo-ET map, ensuring accurate alignment with experimental cryo-ET maps (Fig. 3 stage 3) (SI Table 2). Structural sampling of rigid body positions and orientations as well as flexible bead positions, was performed using the Replica Exchange Gibbs Monte Carlo (MC) algorithm (SI Table 2) (105, 106). Each MC step consisted of a series of random transformations (i.e., rotations and translations) applied to the rigid bodies and flexible beads. The same sampling protocol was used for mBasket modeling, except that the starting structure mimicked the yBasket model (SI Table 3).

### Stage 4: Analysis and validation

Model validation followed four steps (54, 107): (i) selection of the models for validation, (ii) estimation of sampling precision (Fig. S5A), (iii) estimation of model precision, and (iv) quantification of the degree to which a model satisfies the information used and not used to compute it (SI Table 2, 3) (Fig. S5, Fig. S6) (54, 108).

Integrative modeling iterated through the four stages to find a set of models that satisfy our validation criteria listed above. In each iteration, we considered the input information, representation, scoring, and sampling guided by an analysis of models computed in the preceding iteration of the modeling. For example, the initial low precision of the yBasket model encouraged us to improve the resolution of the cryo-ET map by averaging a larger number of subtomograms; and the initial inability to find yBasket models that satisfied both cryo-ET and crosslinking data encouraged us to increase the resolution and flexibility of the coiled-coil representations, re-defining the coiled-coil segments (109), disorder predictions for FG Nups (99), and adding proximity restraints to some components.

**Supplementary Information Table 1.** Information and spatial restraints for Integrative modeling of yeast and mouse basket. For each basket model (column 1), a list of experimental techniques is shown in column 2 that provides the spatial information (column 3) for Integrative modeling. Column 4 provides a reference to the original dataset or method.

| Model | Experimental Technique | Spatial Information | Reference |
| --- | --- | --- | --- |
| Yeast | Quantitative mass spectrometry and biochemical quantification | Stoichiometry of Nups (SI Table 2) | (6) |
|  | Chemical cross-linking with mass spectrometry readout | 626 intra and inter-molecular crosslinks among yNups and yMlps | (6, 24) & this study |
|  | Cryo-ET | The overall shape of the basket including the basket distal density | this study |
|  | Integrative models and atomic structural models | yNup84 complex dimer model yNup2 structural model (SI Table 2) | (5, 110, 111) |
|  | Bioinformatics information | Sequence information coiled-coil propensities of the yMlps alignments. Connectivity between residues. | (109) |
|  | Immuno-EM | Upper and lower bound on the N-/C- terminus of yMlps | (24) |
|  | Biochemical experiments including *in vitro* binding assays proximity labeling | Interactions between yNup60<sup>MBM</sup> (240-318) and yMlp1 (382-620) Interaction between yNup60<sup>N2BM</sup> (505-539) and yNup2 (83-136) nuclear envelope binding domain of yNup1, yNup60 | (26, 55) |
| Mouse/Human | Quantitative mass spectrometry and biochemical quantification | Stoichiometry of Nups (SI Table 3) | (112, 113) |
|  | Cryo-ET | The overall shape of the basket including the basket distal density | this study |
|  | Integrative models and atomic structural models | mNup107 complex dimer model mNup50 structural model (SI Table 3) | (35, 103, 104, 110, 111) |
|  | Bioinformatics information | Sequence information coiled-coil propensities of the mTprs orthologs alignments. Connectivity between residues/beads. Structural equivalent distances between ortholog Nups. | (109) |
|  | Immuno-EM | Upper and lower bound on the N-/C- terminus of mTprs | (102) |
|  | Biochemical experiments including proximity labeling | Nuclear envelope binding domains of mNup153 | (114) |

**Supplementary Information Table 2.**
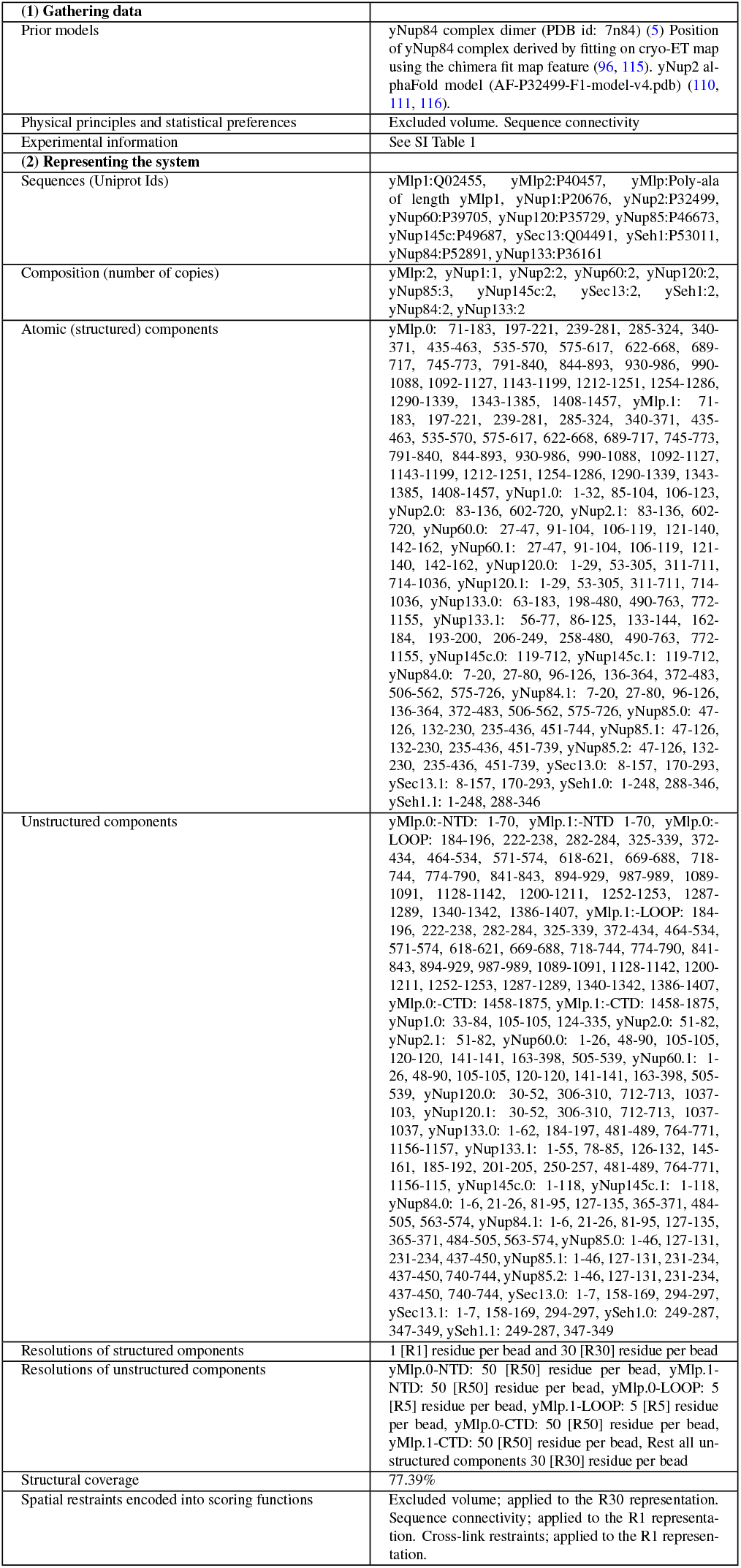
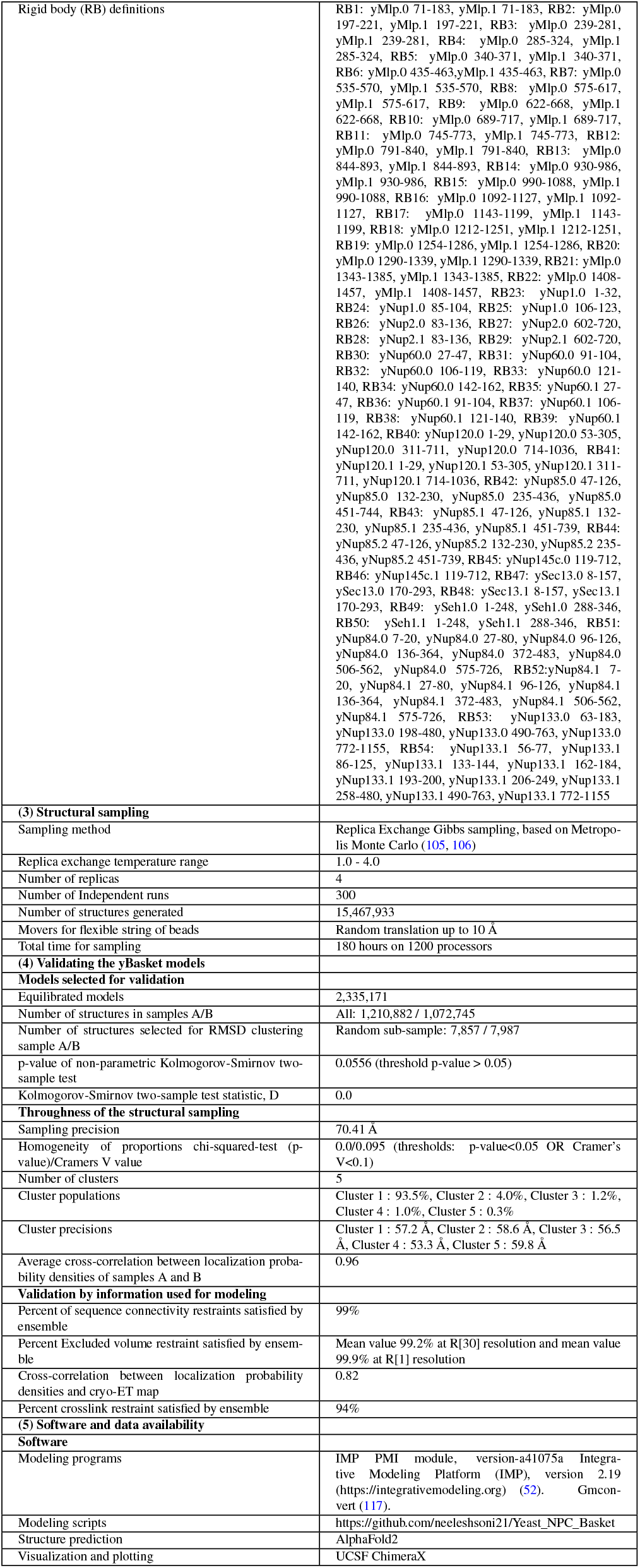
Summary of the Integrative modeling of the yBaskets.

**Supplementary Information Table 3.**
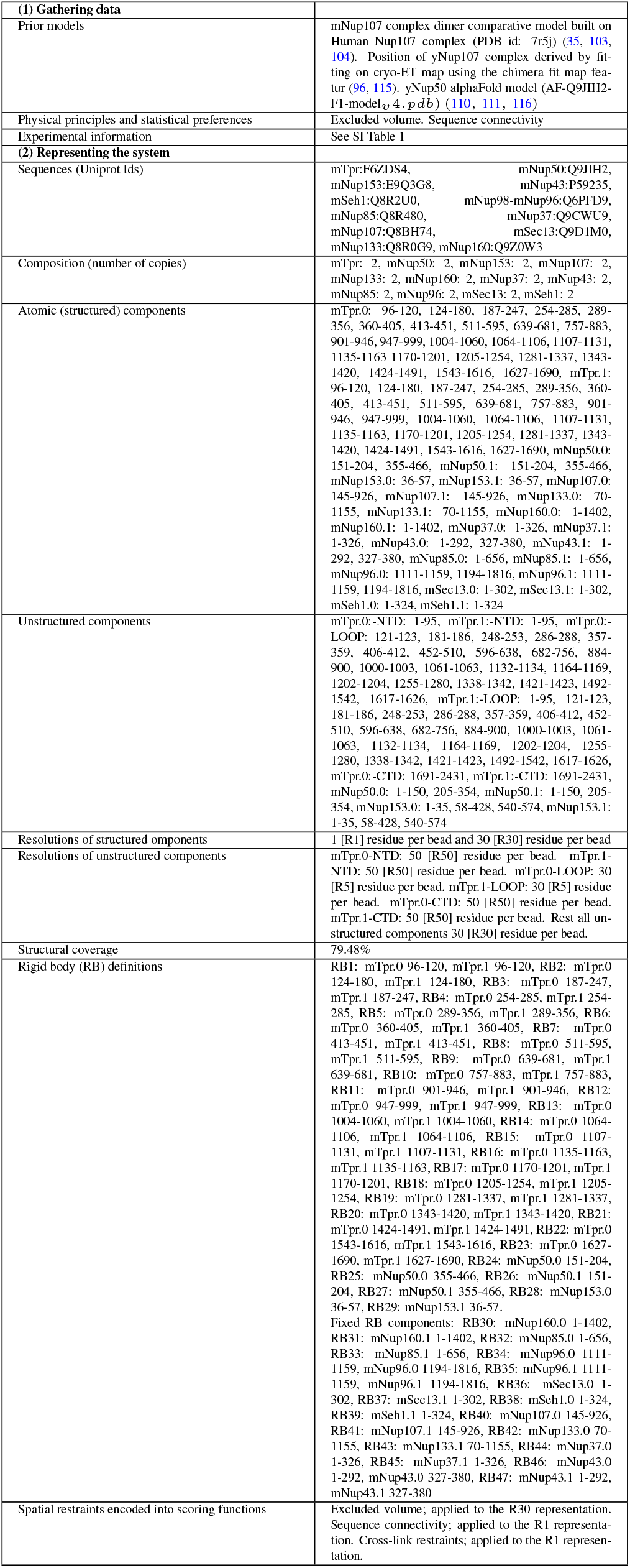
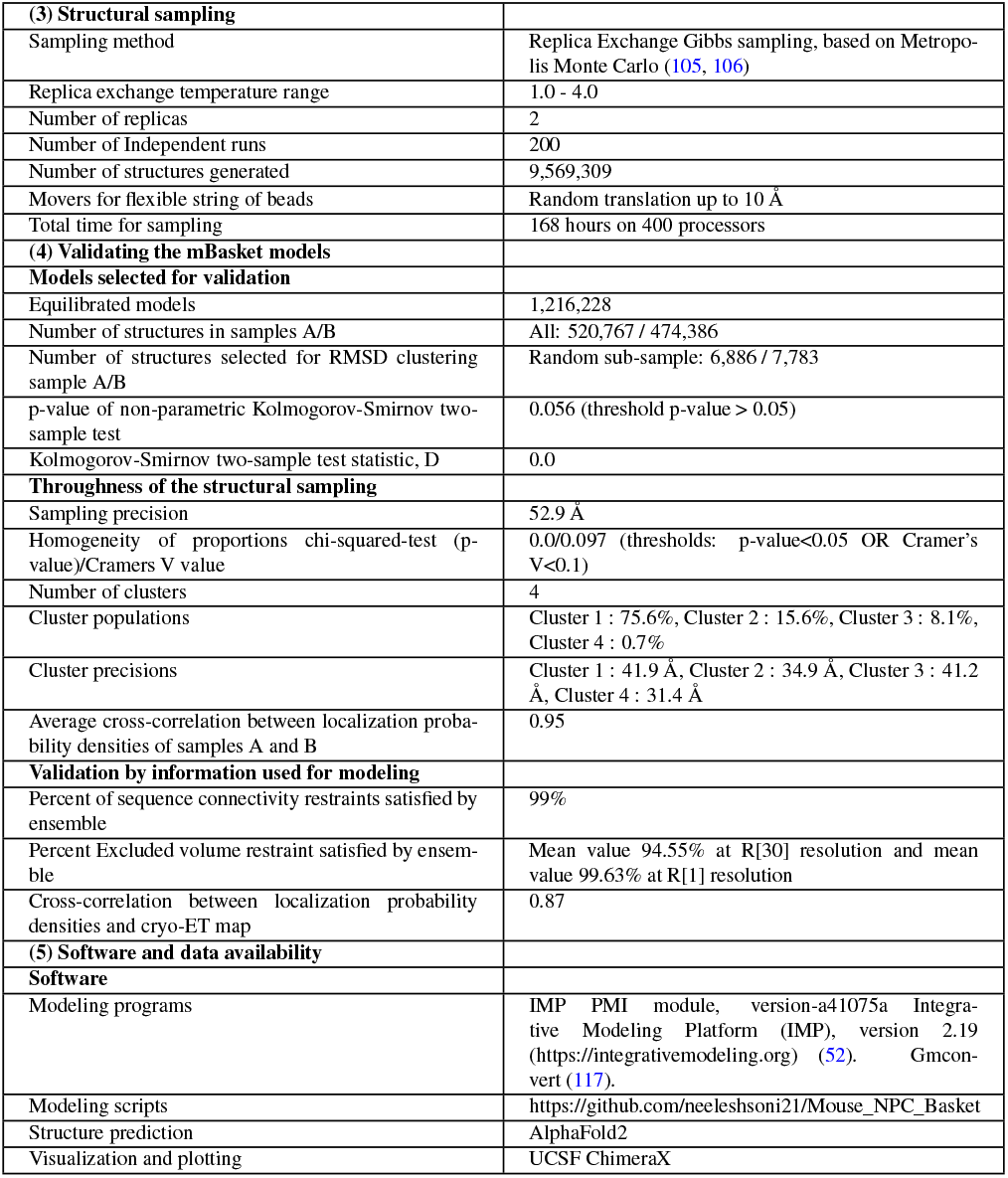
Summary of the Integrative modeling of the mBaskets.

---

**Fig. S1.**
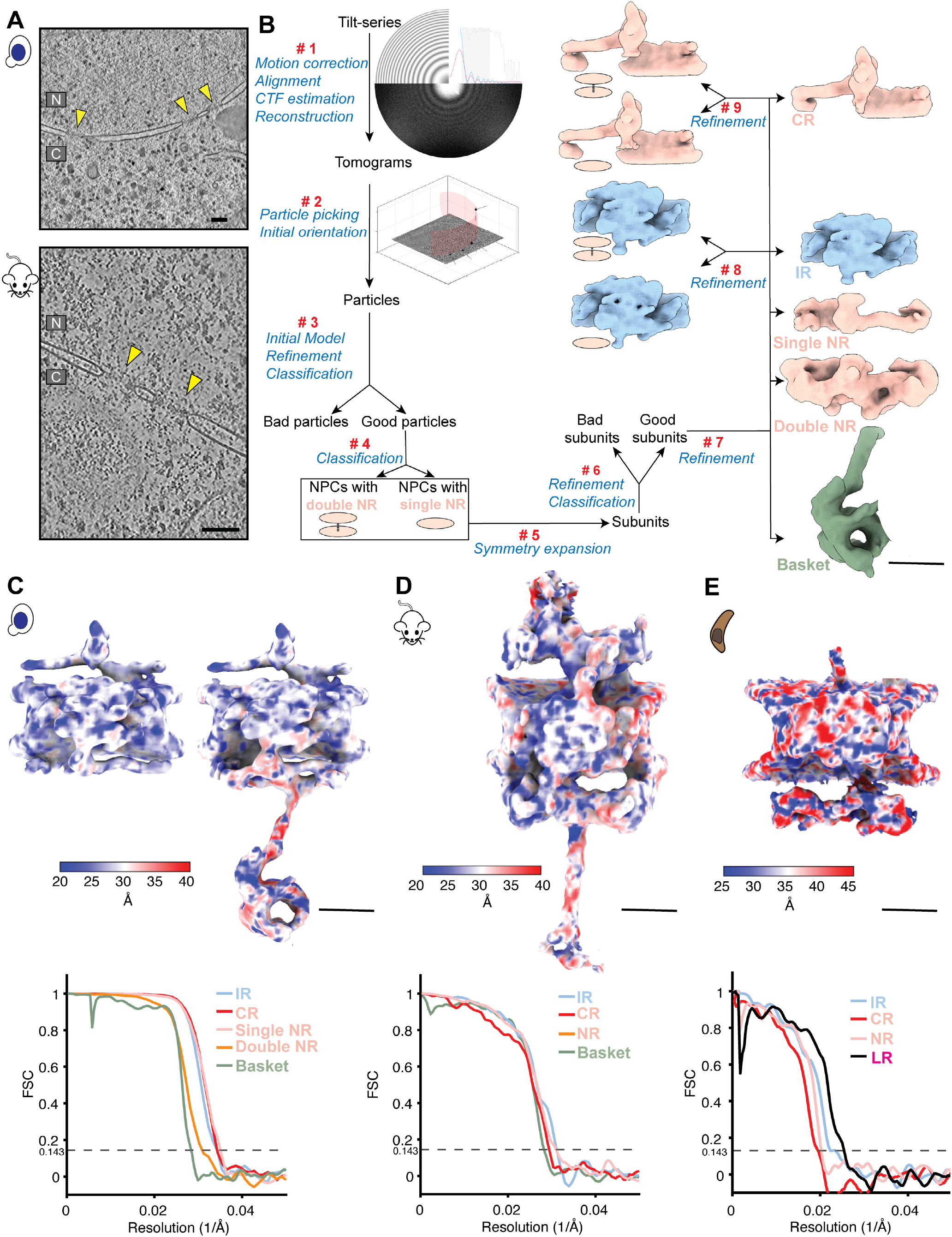
Workflow of the subtomogram analysis of the NPC. (**A**) Slices of representative tomograms from yeast and mammalian cells, where nuclear pores are indicated by yellow arrows in the nucleus. Scale bar: 100 nm. (**B**) Processing workflow of the subtomogram analysis of the NPC, where steps are also indicated in red. The workflow is similar for data from all organisms. Some steps, like 4 for classifying out yNPCs with single and double NR, are more specific for yeast. For yeast, maps of the subunit of IR and CR from NPCs with single and double NR, as shown, resulting from steps 8 and 9 are similar. The same workflow was used for mNPC and pNPC. Scale bar: 200 Å. (C-E) Local resolution maps of the subunit of the NPC (top) along with Fourier shell correlation (FSC) curves of subunits of different rings (bottom) for yeast (**C**), mammalian (**D**), and *T. gondii*. (**E**). Scale bar: 200 Å. CR: Cytoplasmic ring, IR: Inner ring, NR: Nuclear ring, NE: Nuclear envelope, LR: Lumenal ring.

**Fig. S2.**
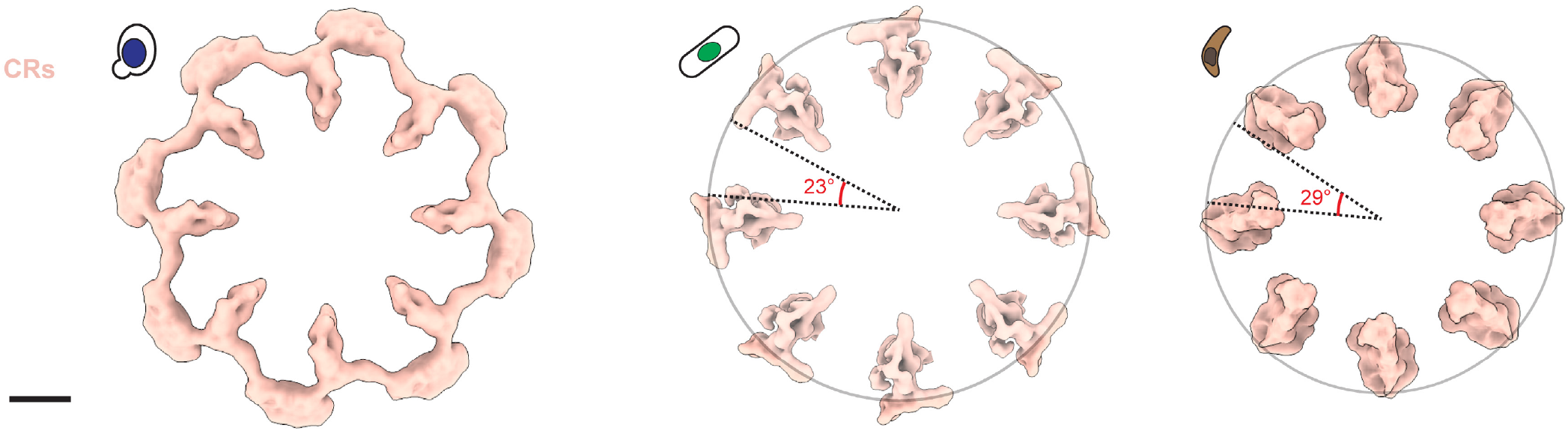
CRs of NPCs from different organisms can have differing extents of completeness. (**A**) The circumference not covered (in central angles), by CRs of *S. cerevisiae* (left), *S. pombe* (middle; EMDB: 11373 (46) and *T. gondii* (right) is 0°(100-0%= 100% covered), 23°× 8 ( 100-51% = 49% covered), 29°× 8 (100-64% = 36% covered), respectively.

**Fig. S3.**
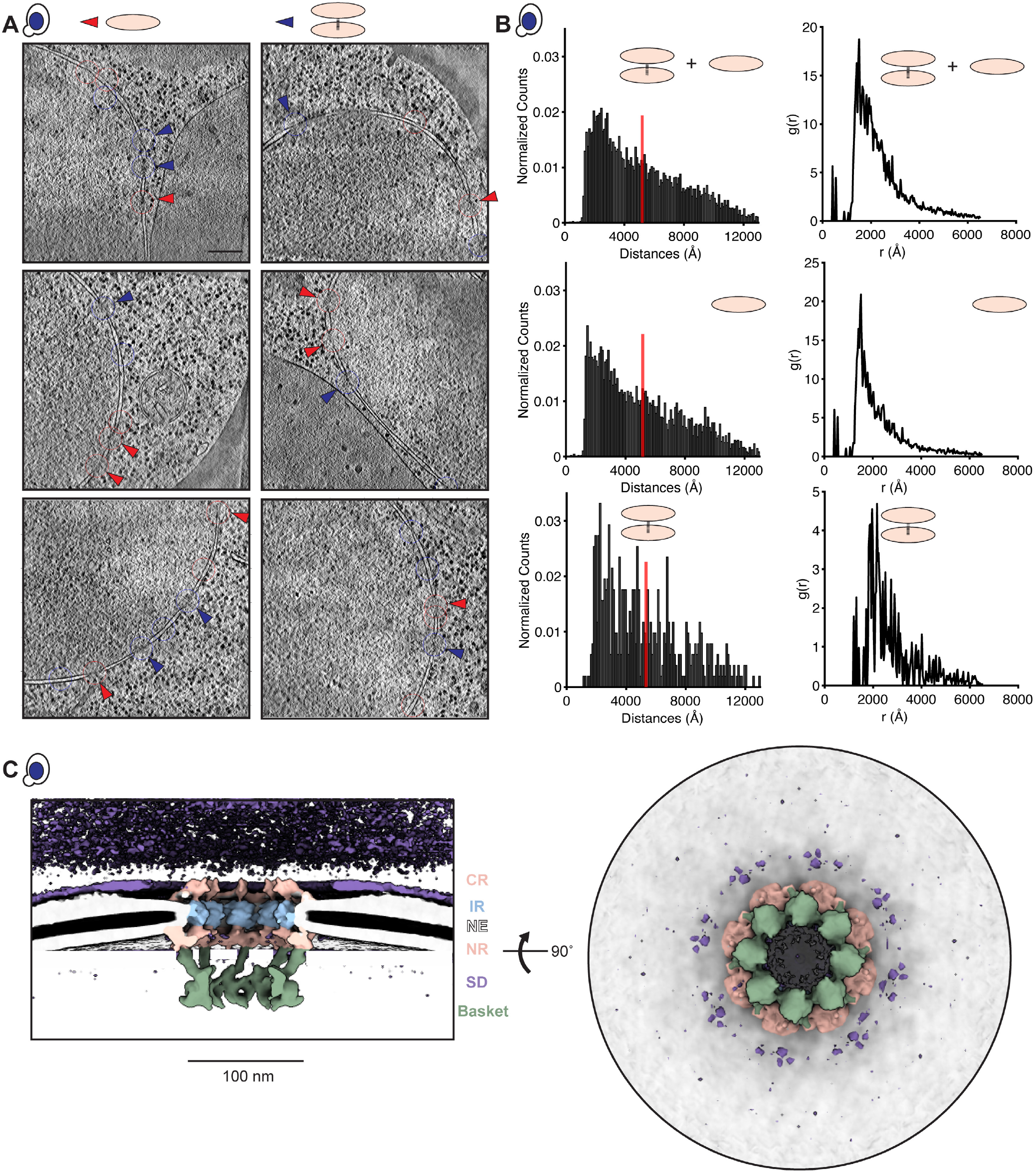
yNPCs have less pronounced nuclear surrounding densities and their single and double NR variants are similarly distributed across the nucleus. (**A**) Slices of representative tomograms from yeast cells highlight yNPCs with single (indicated by red arrows) and double NRs (blue arrows). These images demonstrate that yNPCs with single and double NRs with a stable basket can be adjacent, rather than in distinctly separate areas. Scale bar: 100 nm. (**B**) Analysis of the yNPC’s pairwise distance distribution (left) and the radial distribution function [g(r)] (right), encompassing all NPCs, as well as subsets with either single or double NRs, reveals a similar spatial distribution across the nucleus for all yNPCs. This is evidenced by the qualitative similarity in these distributions. (**C**) The cross-sectional (left) and nucleoplasmic view (right) of the map of yNPC shows the extent of crowding around the yNPC, with the surrounding densities (SD) shown at a very low isosurface threshold. These densities, compared to their counterpart in mNPCs, are much weaker.

**Fig. S4.**
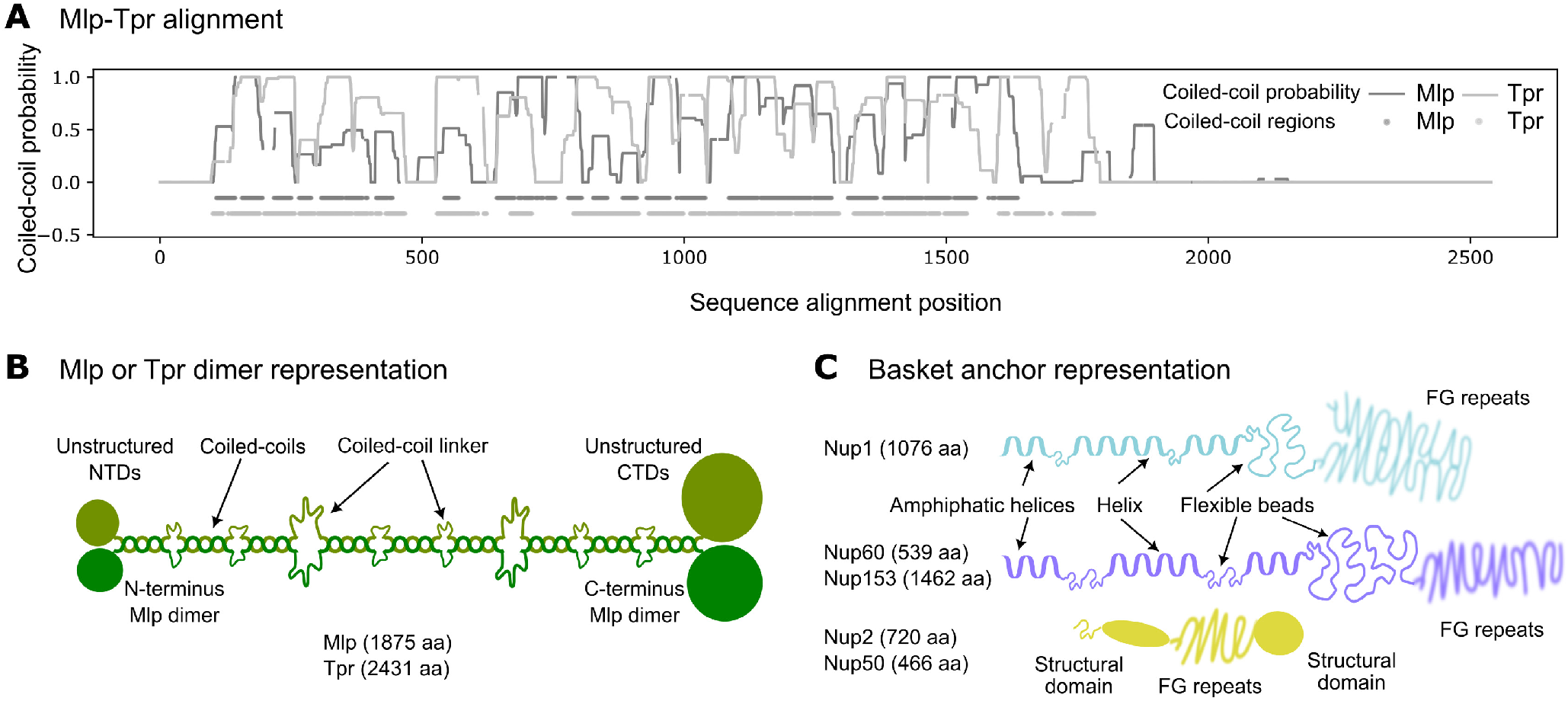
Nuclear basket model representation. (**A**) Coiled-coil probabilities of yMlp (dark gray) and mTpr (light gray) at different positions in their sequence. The region represented as coiled-coil segments is shown as horizontal dots in a straight line below 0.0. (**B**) Schematic representation of a yMlp or mTpr dimer. Coiled-coil regions were represented as rigid bodies and linked through flexible strings of beads. The unstructured N- and C-terminal regions were represented as flexible strings of beads. (**C**) Schematic representation of the FG NR anchor Nups: yNup1, yNup2/mNup50, yNup60/mNup153. All structural regions, including amphipathic helices (AH) at the N terminus, were represented as rigid helices; all non-structural regions, excluding FG repeats, were represented as flexible strings of beads. FG repeats were not included in the model. The number of amino acid (aa) residues for different Nups are indicated in parenthesis.

**Fig. S5.**
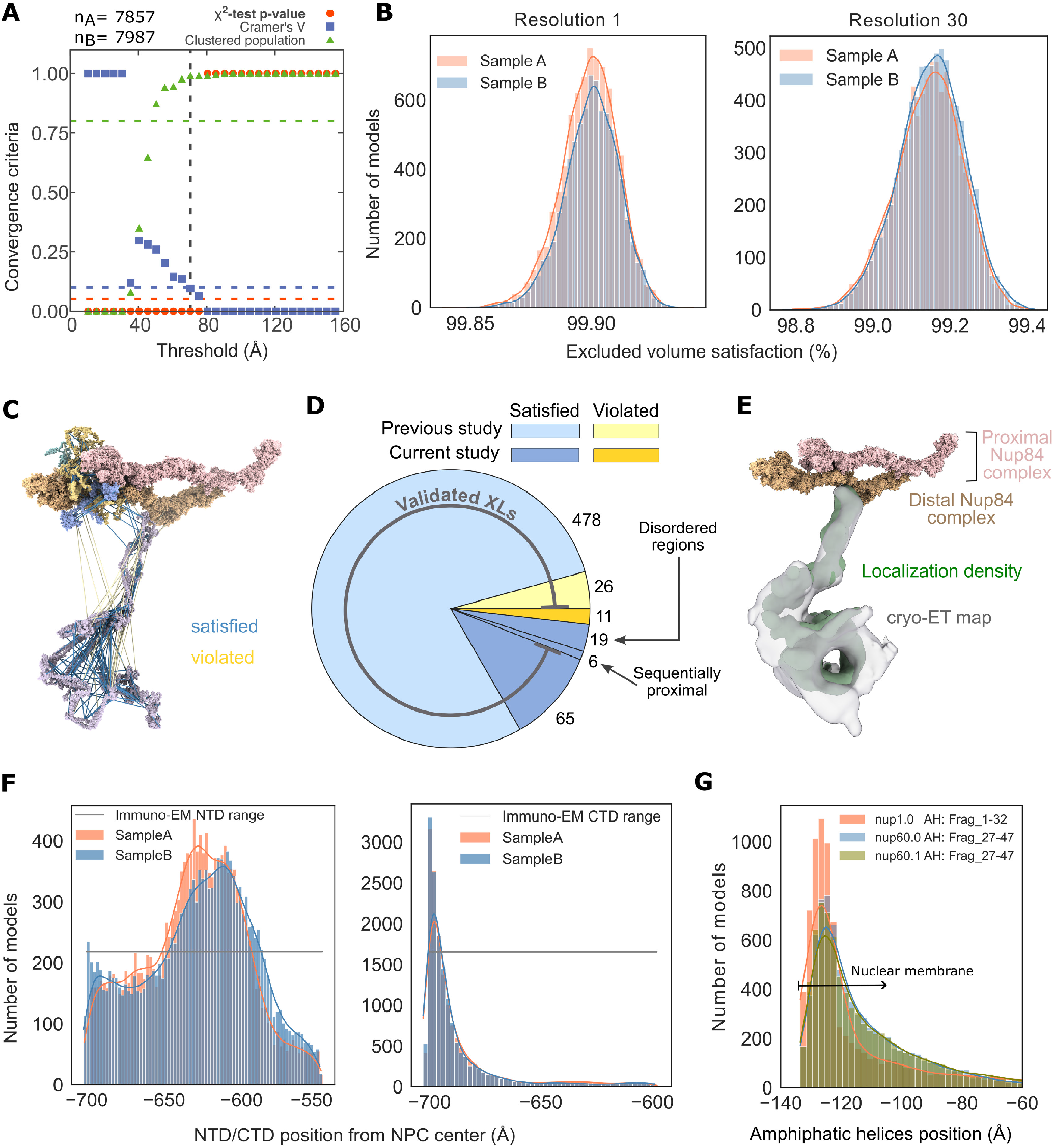
Validation of integrative models of the yBasket. (**A**) Criteria for determining the sampling precision (y-axis), were evaluated as a function of the RMSD clustering threshold (x-axis) (n=15,844 models). First, the P value is computed using the chi-squared-test (one-sided) for homogeneity of proportions (red dots). Second, an effect size for the chi-squared-test is quantified by the Cramers’s V value (blue squares). Third, the population of the structures in sufficiently large clusters is shown as green triangles. The vertical dotted grey line indicates the RMSD clustering threshold at which three conditions are satisfied (Cramer’s V (0.095)<0.1 (blue horizontal dotted lines), P value (0.0)<0.05 and the population of clustered structures (0.99)>0.8 (green horizontal dotted lines, thus defining the sampling precision of 70.4 Å. The three solid curves (red, blue, and green) were drawn through the points to help visualize the results. The number of models in two randomly drawn samples, sample A and sample B, from a pool of filtered models (n=2,283,627 models) is denoted by nA (7857 models) and nB (7987 models). (**B**) Histogram of excluded volume restraint satisfaction of samples nA and nB at two different resolutions of 1 and 30 residues per bead. (**C**) Chemical cross-links were mapped onto the structure of the yBasket subunit with lines. Satisfied crosslinks where Euclidian Ca–Ca distances below 35 Å in the model ensemble are represented with blue lines, whereas violated crosslinks are with red lines. (**D**) A pie chart of satisfied (shades of blue) and violated (shades of yellow) crosslinks grouped into previously published (6) (light blue and light yellow) and current study (dark blue and dark yellow) crosslinks. The validated crosslinks (gray arc) account for 94% of the crosslinks; validated crosslinks have MS2 spectra with multiple backbone fragmentations of both peptides and peptides of at least 6 residues. Of the non-validated crosslinks, 11 were violated, 6 were trivially satisfied due to sequence proximity, and 19 were in the disordered/N-terminus/C-terminus regions of the basket distal density of our model. (**E**) A side view of the yBasket models localization density attached to the distal yNup84 complex spans most of the cryo-ET density map. (**F**) Satisfaction of experimentally derived range (gray line) of the position of yMlps N- and C-terminus by the model ensemble derived values (coral and light blue). (**G**) Satisfaction of experimentally derived range (gray line) of the position of amphipathic transmembrane helices of yNup1 and yNup60 by the model ensemble derived values (coral and light blue).

**Fig. S6.**
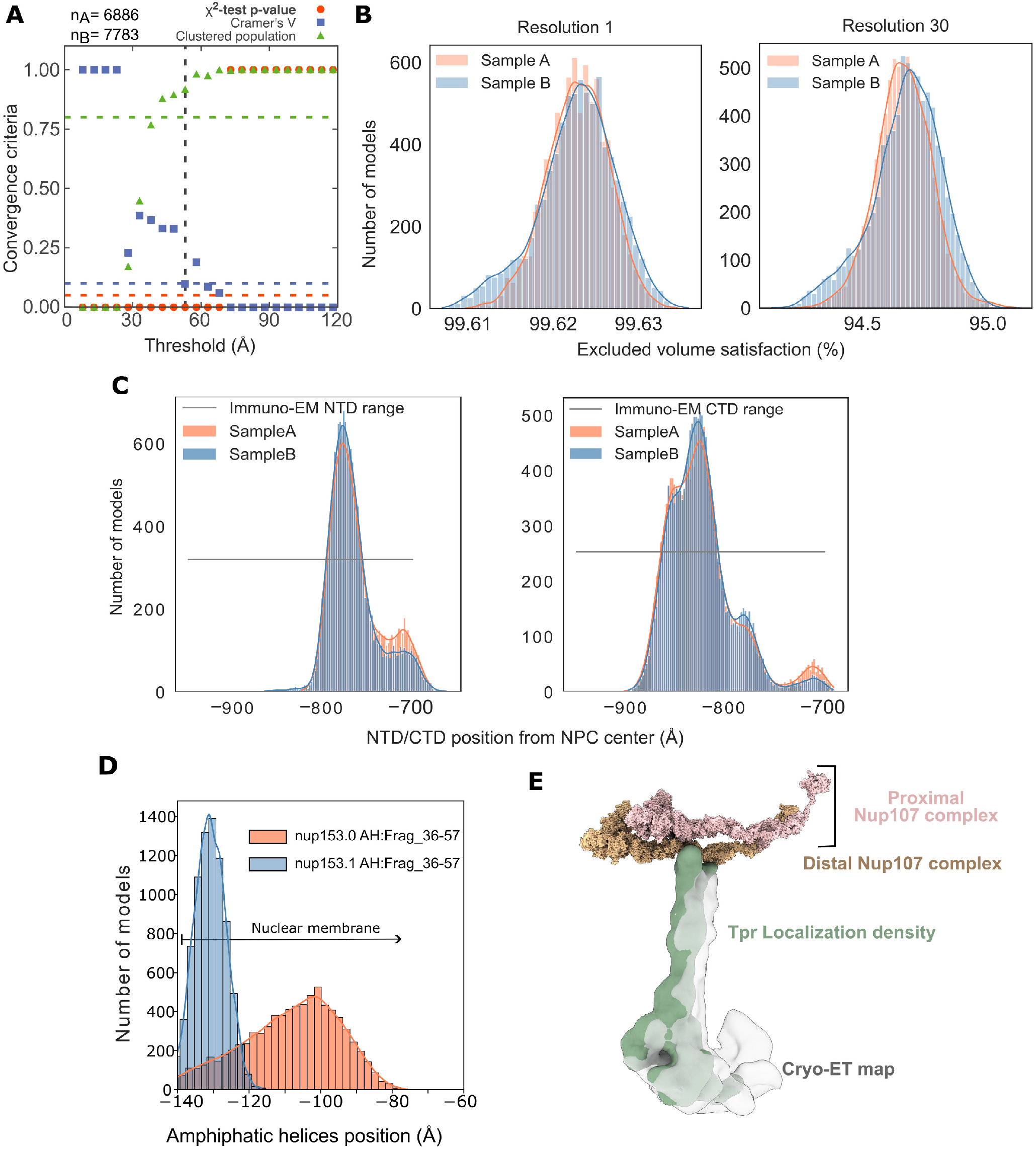
Validation of integrative models of the mBasket. (**A**) Criteria for determining the sampling precision (y-axis), were evaluated as a function of the RMSD clustering threshold (x-axis) (n=14,669 models). First, the P value is computed using the chi-squared-test (one-sided) for homogeneity of proportions (red dots). Second, an effect size for the chi-squared-test is quantified by the Cramers’s V value (blue squares). Third, the population of the structures in sufficiently large clusters is shown as green triangles. The vertical dotted grey line indicates the RMSD clustering threshold at which three conditions are satisfied (Cramer’s V (0.097)<0.1 (blue horizontal dotted lines), P value (0.0)<0.05 and the population of clustered structures (0.92)>0.8 (green horizontal dotted lines, thus defining the sampling precision of 52.9 Å. The three solid curves (red, blue, and green) were drawn through the points to help visualize the results. The number of models in two randomly drawn samples from a pool of filtered models (n=995,151 models) is denoted by nA (6886 models) and nB (7783 models). (**B**) Histogram of excluded volume restraint satisfaction of samples nA and nB at two different resolutions of 1 and 30 residues per bead. (**C**) Satisfaction of experimentally derived range (gray line) of the position of mTpr’s N- and C-terminus by the model ensemble derived values (coral and light blue). (**D**) Satisfaction of the experimentally derived range (gray line) of the position of amphipathic transmembrane helices of mNup153 by the model ensemble derived values (coral and light blue). (**E**) A side view of the mBasket models localization density attached to the distal mNup107 complex spans most of the cryo-ET density map.

